# Endogenous circadian reporter cell lines as an efficient platform for studying circadian mechanisms

**DOI:** 10.1101/2022.06.23.497383

**Authors:** Jiyoung Park, Kwangjun Lee, Hyeongseok Kim, Heungsop Shin, Choogon Lee

## Abstract

Adverse consequences from having a faulty circadian clock include compromised sleep quality and poor performance in the short-term, and metabolic diseases and cancer in the long-term. However, our understanding of circadian disorders is limited by the incompleteness of our molecular models and our dearth of defined mutant models. Because it would be prohibitively expensive to develop live animal models to study the full range of complicated clock mechanisms, we developed *Per1-luc* and *Per2-luc* endogenous circadian reporters in a validated clock cell model, U2OS, where the genome can be easily manipulated, and functional consequences of mutations can be accurately studied. Using these reporter cells, we uncovered critical differences between two paralogs of *Per* and *Cry*, as well as working principles of the circadian phosphotimer. Our system can be used as an efficient platform to study circadian sleep disorders such as Familial Advanced Sleep Phase Syndrome (FASPS) and their underlying molecular mechanisms.

## Introduction

The circadian clock drives daily rhythms in behavior and physiology (Bell-Pedersen et al., 2005; Harmer et al., 2001; Mohawk et al., 2012; Reppert and Weaver, 2002; Schibler, 2005), and dysfunction or disruption of the clock has been implicated in diverse disease states including sleep disorders (Bass and Takahashi, 2010; Drake et al., 2004; Green et al., 2008; Musiek et al., 2013; Zhu and Zee, 2012). Decades of prior work have revealed that the clock is built on a core transcriptional feedback loop that is cell autonomous, involving transcriptional and post-translational regulation of the pacemaker *Period* (*Per*) genes (Patke et al., 2020; Potter et al., 2016). In the feedback loop, the activator complex CLOCK:BMAL1 drives transcription of the pacemaker genes *Per1* and *Per2* (*Per*) along with many other clock-controlled genes. PER proteins form an inhibitory complex that also contains CRYPTOCHROME (CRY) proteins and casein kinase CK1δ/ε proteins. The circadian phase, e.g., onset of activity or sleep, is determined by the oscillations of this PER/CRY/CK1 complex (Chen et al., 2009; Lowrey et al., 2000; Toh et al., 2001).

The paralogs of *Per* and *Cry* genes share some common/redundant functions but also differ in function and regulation. In the case of *Per*, the paralogs *Per1* and *Per2* (*Per3* is considered nonessential) seem to play redundant roles in the master clock tissue, the SCN, as knockouts of either gene produce little period and phase alterations in behavioral rhythms (Pendergast et al., 2010; Shearman et al., 2000). However, the regulation of *Per1* and *Per2* may differ dramatically in peripheral tissues and cell culture. For example, *Per1* transcription in cultured cells is rapidly induced by high serum and forskolin, but *Per2* transcription is marginally affected by these signals (Balsalobre et al., 2000). In mouse tissues, phases of *Per1* transcript and protein rhythms are advanced relative to those of *Per2* (Lee et al., 2001).

Therefore, the circadian period and phase of PER protein oscillations in *Per1* KO cells and *Per2* KO cells may differ significantly, likely because of different kinetics in mRNA and protein profiles in these peripheral cells. This is an unexplored critical issue because circadian properties such as phase of all clock-controlled genes (ccgs) can be affected when only one *Per* is present compared to wt cells. In the case of *Cry,* knockouts (KOs) of *Cry1*, *Cry2,* and *Cry1/2* exhibit shortened rhythms, lengthened rhythms, and arrhythmicity, respectively. Although it has been suggested that difference in binding affinity of two CRYs to CLOCK:BMAL1 is the molecular underpinning for the opposite period alteration in *Cry* single KOs (Fribourgh et al., 2020; Rosensweig et al., 2018), it has not been studied how two CRYs differentially affect the pacemaker genes *Per1* and *Per2*. Period alteration in *Cry* KOs would be manifested ultimately through altered regulation of *Per* at transcriptional and posttranscriptional levels.

Another critical issue in the field is how PER phosphorylation by CK1δ and CK1ε can be extended to over ∼12 hours, which is responsible for the prolonged circadian feedback loop. This is an unconventional kinase-phosphorylation relationship, especially considering that PER and CK1 make an unusually stable interaction through a dedicated domain in PER called CKBD (Casein Kinase Binding Domain) (Lee et al., 2001; Lee et al., 2004; Vielhaber et al., 2000). A series of phosphorylation steps across several serine residues in this domain seems to play an important role in tuning phase and period (Fribourgh et al., 2020; Narasimamurthy et al., 2018; Philpott et al., 2020). The Ser662Gly mutation in the CKBD of *Per2* gene causes advanced phase and shortened period in humans, leading to a condition called Familial Advanced Sleep Phase Syndrome (FASPS) (Toh et al., 2001). Although the FASPS motif has been a main focus of studies of the CKBD due to its defined role in the human sleep disorder, CKBD has other conserved motifs, and it has not been studied how these motifs contribute to the phosphorylation-mediated timing system.

Because the circadian clock is cell autonomous, genetic disruptions of the clock manifest similar phenotypes at the behavioral and cellular levels, and cell culture has proven to be a valuable and valid platform for characterizing the molecular biology of circadian rhythms (Chen et al., 2009; Liu et al., 2007; Yoo et al., 2004). The endogenous clocks of cultured cells— including mouse embryonic fibroblasts (MEFs) and human U2OS cells—can be precisely measured in real time by introducing a luciferase (Luc) reporter gene under control of a clock promoter (Liu et al., 2007; Maier et al., 2009; Yamazaki et al., 2000; Yoo et al., 2004). Across numerous studies, such cells have served as functional models for in vivo circadian clocks, and results have been consistently validated in live animal models. Cell culture models are not only less resource-consuming, but also more easily manipulated by chemicals and transgenes, which makes the cell models more suitable for mechanistic studies. One key limitation in cell models today is the lack of endogenous phase and period reporters other than the *mPer2-Luc* reporter in MEFs (Yoo et al., 2004), which is a mouse, not a human cell model. Exogenously transfected reporters like *Per2-* or *Bmal1*-promoter controlled Luc are useful to an extent but do not accurately reflect the endogenous status of the clock.

In this study, we generated human *Per1-luc* and *Per2-luc* endogenous knockin genes in U2OS cells, to produce a human cell model with robust bioluminescence rhythms. When knockout phenotypes for major clock genes were quantified by bioluminescence rhythms in these cells, they were consistent with phenotypes of knockout mice. We further used these cells to uncover critical differences between the paralogs of *Per* and *Cry*, and to gain critical insights into how timing cues are precisely generated by the phosphotimer through dynamic interaction between PER and CK1δ/ε.

## Results

### Generation of U2OS cells with endogenous *Per1-luc* and *Per2-luc* knockin genes and robust rhythms in bioluminescence and clock proteins

The main challenge in developing reporter knockins (KIs) was that the screening process could be very cumbersome due to a high number of off-target insertions (Uddin et al., 2020; Zhang et al., 2015). We have tested several established protocols as well as newer ones employing high fidelity CAS9 (HF CAS9) and the paired Cas9D10A nickase approach to decrease off-target insertions (Kleinstiver et al., 2016; Koch et al., 2018; Slaymaker et al., 2016), but they all produced different off-target insertions and offered no meaningful advantage over wtCAS9 for on-target KI generation. To streamline the selection process for positive clones after single-cell sorting, we employed a dual reporter system that eliminates the need for molecular experiments during the selection process (Figure 1A). To generate *Per1* and *Per2* reporter lines, U2OS cells were transfected with two plasmids, all-in-one GFP-sgRNA-CAS9 (Ran et al., 2013) and repair templates with luciferase-T2A- mRuby3 (Figure 1A). If targeting is successful, Luc-T2A-mRuby3 will be inserted between the last amino acid (AA) and stop codon in each *Per* gene (Figure 1A). The T2A cleavage site was added because Luc alone has been proven not to disrupt PER function or clock function in the *mPer2^Luc^* KI mouse (Yoo et al., 2004) whereas this has not been proven for the bulkier Luc- mRuby3 dual tag. In the first selection after transfection, stable mRuby3-expressing cells were singly sorted into 96 well plates by FACS. Because we hypothesized mRuby3 expression would be low based on prior work with the *mPer2-GFP* KI mouse (Smyllie et al., 2016), we selected a low range of red signal (Figure 1B) for single cell sorting; this tunability is a major advantage of FACS versus antibiotic selection. Subsequent bioluminescence selection allowed us to select clones with robust rhythms without the need for expansion, genomic DNA prep and PCR analysis from individual clones (Figure 1C). We isolated 7 final clones for *Per1* (4 hetero- and 3 homozygous KIs), and 6 final clones for *Per2* (all heterozygotes). The 2^nd^ selection produced ∼30% rhythmic clones. These final clones were further validated by junction PCR and sequencing (Figure 1D and S1), and by immunoblotting with anti-PER and Luc antibodies (Figure 1E and S2). Finally, single specific insertions were verified by showing a complete loss of bioluminescence and mRuby3 signals when frame-shifting mutations were introduced in early exons in *Per* genes (Figure S3).

**Figure 1.**
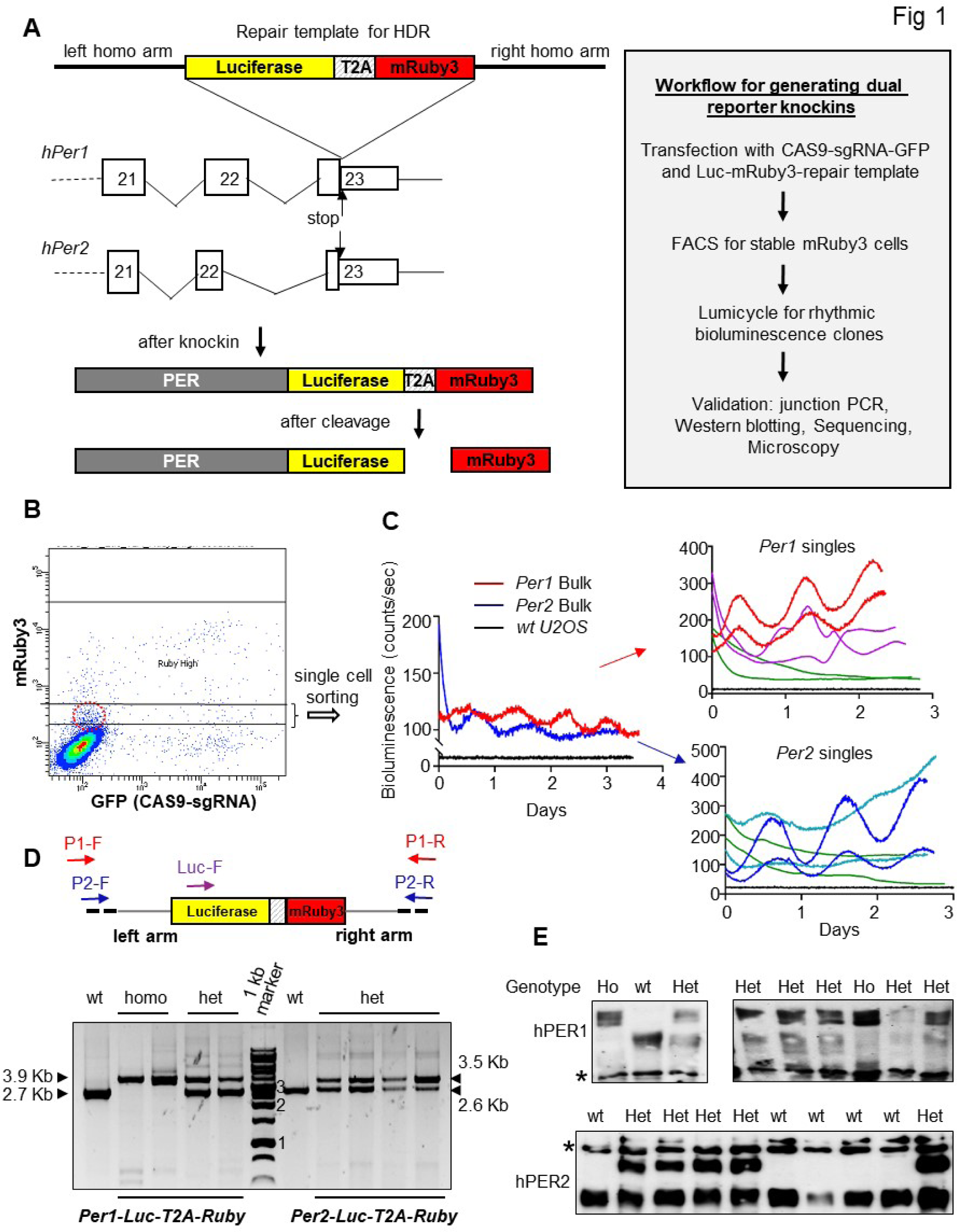
A streamlined selection process using a tandem reporter system successfully and rapidly identified knockin (KI) reporter cells. (A) KI schematics using a tandem reporter, fluorescence plus bioluminescence. In heterozygote KI clones, the non-KI alleles had minor mutations near the stop codon due to indels without KI (Figure S1). (B) Cells expressing mRuby3 in a low range were selected in bulk and singly sorted into 96-well plates by FACS. (C) The bulk sorted cells and single clones were set up in Lumicycle 96. Note that red and blue traces represent specific *Per1^Luc^* and *Per2^Luc^* KI clones, respectively. (D) Three-primer PCR using two primers binding outside of the homologous arms and a primer binding to luciferase was done to screen for on-target KI. (E) On-target KI (Ho = homozygous, Het = heterozygous) was confirmed by immunoblotting using anti-PER and anti-Luc antibodies (Fig S2). * non- specific band. See also Figures S1-S3.

We confirmed that the KIs did not cause any disruption in clock function by comparing the molecular rhythms between PER and PER-Luc proteins in heterozygote clones (Figure 2). When comparing *Per1* and *Per2* heterozygote KI clones (hereafter referred to as *Per1^Luc^* and *Per2^Luc^*) (Figure 2A), PER1-Luc peaked 2-3 hrs earlier than PER2-Luc, which is consistent with the phases of PER rhythms in mouse peripheral tissues (Lee et al., 2001). PER1-Luc signals were higher than those of PER2-Luc reporters (Figure 2A, S4A and B). When these *Per^Luc^*reporters were compared to the transgenic *Bmal1-Luc* reporter (Liu et al., 2007), they showed almost antiphase oscillations, consistent with the natural antiphase oscillations of *Per* vs *Bmal1* mRNA in vivo (Figure 2B). The periods of *Per1* and *Per2* KI reporters were not different from each other (Figure 2C and S4C), but they were slightly different from that of the transgenic *Bmal1- Luc* reporter. The mouse endogenous *mPer2^Luc^* reporter in MEFs produced similarly antiphase oscillations compared to the *Bmal1-Luc* reporter in U2OS cells (Figure 2D). Both PER1-Luc and PER2-Luc fusion proteins showed similarly robust oscillations in abundance and phosphorylation as their wt counterparts in the heterozygote clones (Figure 2E and 2F). Robust oscillations of PER-Luc fusion proteins were also confirmed by immunoblotting with anti-Luc antibody (Figure 2E-G). Finally, their functionality was verified by confirming that PER-Luc- containing clock complexes exhibited the same abundance and phosphorylation rhythms as their wt PER-containing counterparts (Figure 3A).

**Figure 2.**
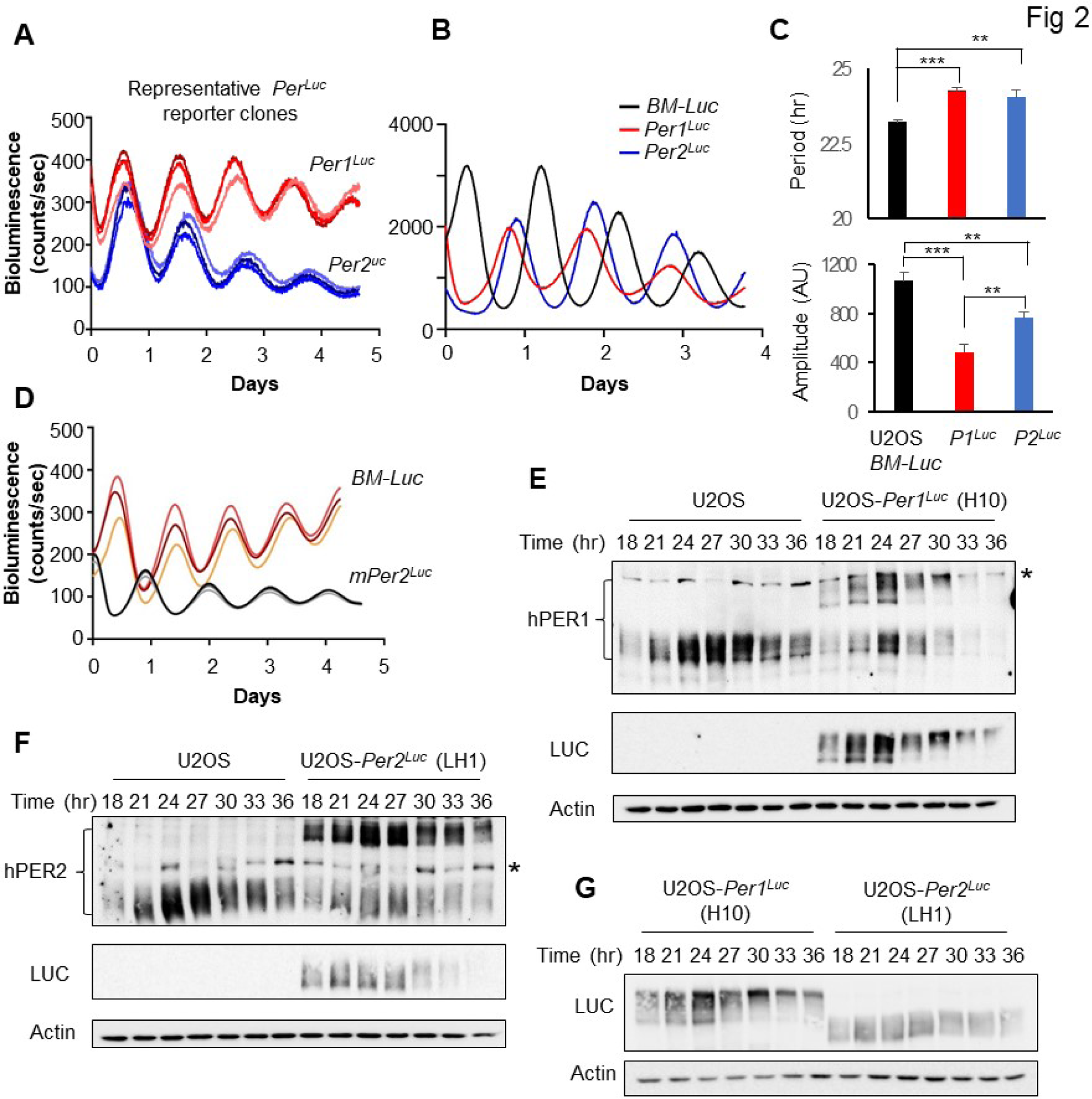
Endogenous *Per^Luc^* reporters exhibit robust oscillations without disrupting the clock mechanisms. (A) Bioluminescence rhythms are shown for representative heterozygous *Per1^Luc^* and *Per2^Luc^*clones, using a similar number of cells. Note that PER1-Luc peaks earlier than PER2-Luc. Three replicates are shown for each clone. Homozygous *Per1^Luc^* clones showed higher signals than heterozygous clones demonstrating that reporter cells are highly quantitative (Fig S4A). More clones are shown in Fig S4B, C. (B) *Per1^Luc^*and *Per2^Luc^* exhibited antiphase oscillations to transgenic *Bmal1* promoter-Luc reporter in U2OS cells. (C) Circadian period of *Per1^Luc^* and *Per2^Luc^* reporters is not significantly different. Representative of three experiments. Mean+/-SD. *<0.05; **<0.01; ***<0.001. (D) m*Per2^Luc^* MEFs showed antiphase oscillations to *Bmal1-Luc* U2OS cells. (E-G) PER-Luc fusion proteins showed robust oscillations in abundance and phosphorylation almost identical to their wt counterparts. See also Figure S4.

**Fig 3.**
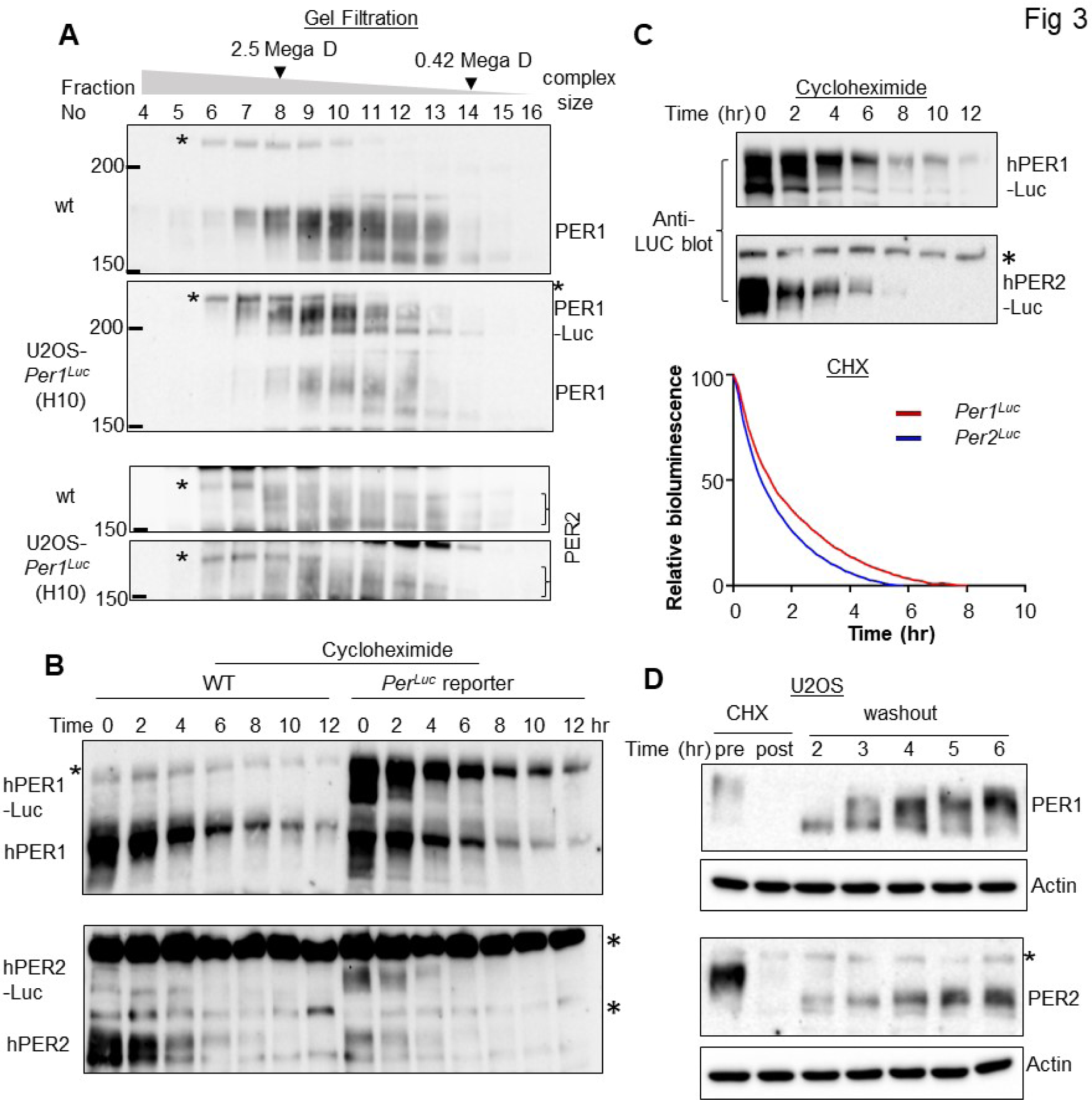
PER1 and PER2 are significantly different in key parameters. (A) Reporter KI did not disrupt endogenous complex formation. * non-specific band. Representative of two experiments. (B, C) PER1 was significantly more stable than PER2. Representative of three experiments. Mean of three traces are shown. p<0.001. (D) PER1 phosphorylation and accumulation were significantly faster than those of PER2 (p<0.001, see Fig S5A for quantitative analysis). Existing PER was depleted by CHX treatment for 8 hrs (post). See also Figure S5.

We have previously shown that the level of PER phosphorylation highly correlates with the size of clock protein complexes because PER hyperphosphorylation is induced by multimerization of PER monomers (Beesley et al., 2020). Studying our novel PER reporter cells uncovered interesting differences between PER1 and PER2 at the posttranslational level.

Although both PER1 and PER2 are much less stable than other clock proteins (D’Alessandro et al., 2017), PER2 was significantly more unstable than PER1 (Figure 3B and 3C). This was verified by immunoblotting of time course samples and real-time bioluminescence (Figure 3C). More significantly, regarding the phosphotimer function of PER, PER1 phosphorylation occurs at a much higher rate than that of PER2 (Figure 3D). PER1 accumulated faster than PER2 after existing proteins were depleted by a long cycloheximide treatment because PER1 was more stable (Figure 3D and S5A). These data have important implications when either *Per1* or *Per2* is inactivated because although either one can sustain the clock as the sole pacemaker, the resulting oscillations could be very different due to their differences in these key parameters.

### Phase of circadian genes is reversed in *Per1* knockout cells

Numerous lines of evidence indicate that the phases of circadian behaviors such as wake and sleep timing (activity onset and offset) are determined by phases of PER oscillations. For example, new wake/sleep cycles following transmeridian travel are established by altered phases of *Per* genes through re-aligning of *Per* oscillations to the altered light cycles (Reppert and Weaver, 2001, 2002; Shigeyoshi et al., 1997). *Per* genes are the only clock genes that can be phase-shifted directly by light in the SCN. We measured how period and phase of *Per1* and *Per2* genes are affected in non-light-responsive peripheral clock cells (U2OS) by examining PER-Luc rhythms when the other paralog is absent. When *Per2* was knocked out in *Per1^Luc^* cells (Figure 4A, 4B and 4D), the period and phase of PER1-Luc rhythms were little changed. However, when *Per1* was knocked out in *Per2^Luc^* cells (Figure 4A, 4C and 4D), the phase of PER2-Luc, but not the period, was dramatically altered, almost resulting in phase reversal. A similarly little altered and dramatic phase shift in *Per1* KO and *Per2* KO U2OS cells, respectively, were observed with wt *Per* paralogs using the transgenic *Bmal1-Luc* reporter (Figure 4E and 4F), showing that the phase shift was not an artifact of the PER-Luc fusion. The dramatically altered phase in PER2 rhythms in *Per1* KO cells was also confirmed by immunoblotting (Figure 4G). This phase reversal is not a unique response to the use of 50% horse serum (‘serum shock’) as the phase-setting stimulus (zeitgeber): similar responses of *Per1* and *Per2* KOs were observed by a different zeitgeber, forskolin (Figure S5B). These data strongly suggest that the phase of all circadian genes, not just core clock genes, are phase-reversed in peripheral clock cells when *Per1* is deleted or inactivated. These data also suggest that *Per1* phase is predominant over *Per2 in* wt cells, and the resetting mechanism for *Per2* is very different from that of *Per1* as suggested above (Figure 2 and 3). Our data are consistent with previous studies by the Schibler group showing that immediate transcriptional response to zeitgebers is very different between *Per1* and *Per2* genes (Balsalobre et al., 1998; Balsalobre et al., 2000). As with mRNA levels during early hours after 2 hr serum shock (Balsalobre et al., 2000), PER1 protein increased more dramatically than PER2, but their phases were similar in wt cells (Figure 4H). However, in *Per1* KO cells, PER2 profile in abundance and phosphorylation was very different from that in wt cells, further supporting that the resetting mechanism for *Per2* by zeitgebers is very different when *Per1* is present versus when it is absent.

**Fig 4.**
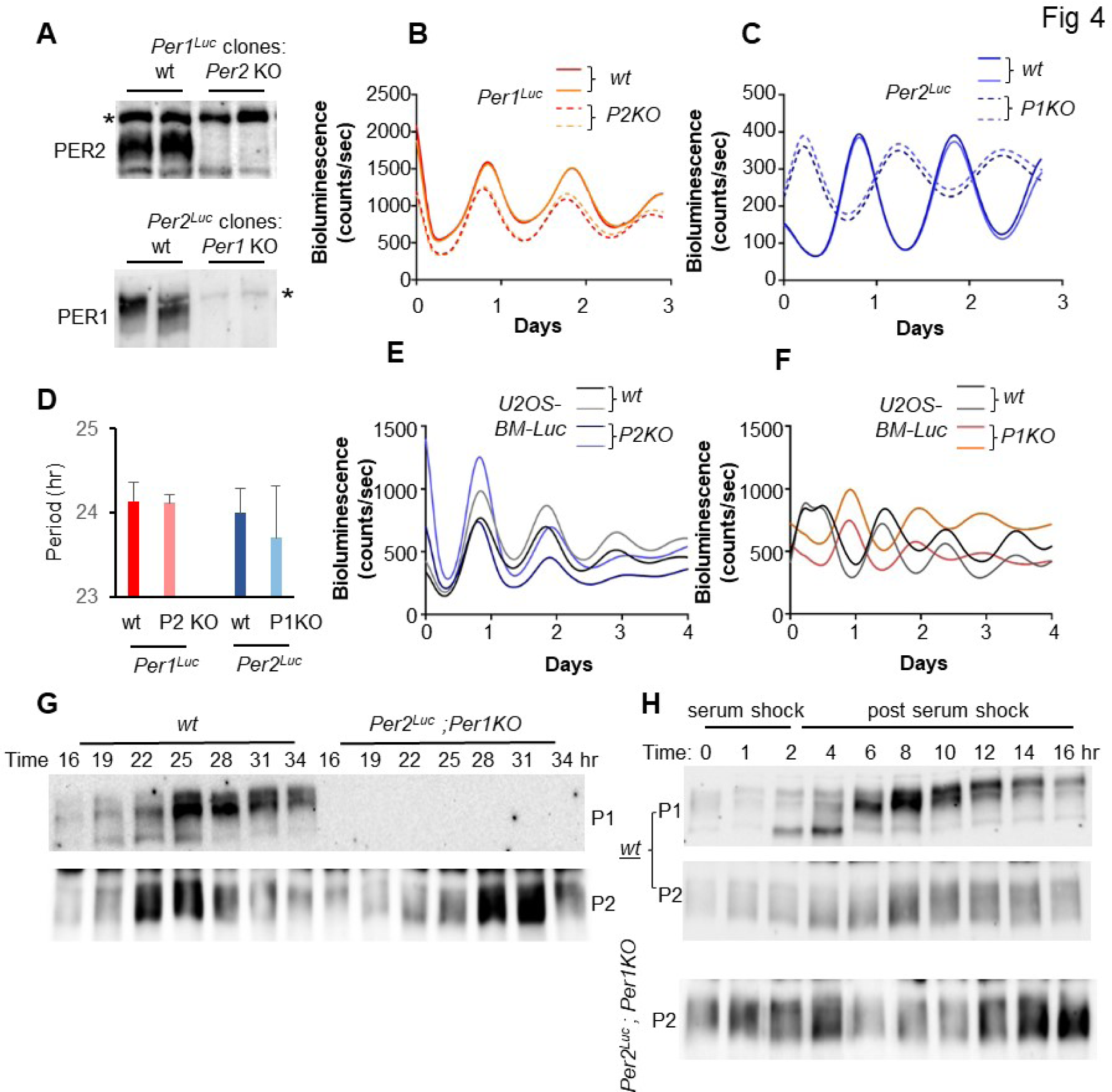
*Per2* phase is reversed in the absence of the *Per1* gene in peripheral cells. (A) *Per* KO was screened by immunoblotting. (B, C) Bioluminescence rhythms of representative *Per2* KO in *Per1^Luc^* and *Per1* KO in *Per2^Luc^* cells are shown. Two traces are shown for each clone. N=3. (D) Circadian period was not significantly different between *Per* KO and wt reporter cells. Mean+/- SD. N=3. Representative of three experiments. (E, F) *Per* KO in *Bmal1-Luc* U2OS cells showed similar results. Two traces are shown for each clone. (G) Antiphase oscillations of PER2 between wt and *Per1* KO cells were confirmed by immunoblotting for PER2. (H) Early response of PER2 to horse serum treatment (an established zeitgeber for cultured cells) was dramatically different between wt and *Per1* KO cells. Cells were given a 2-hour serum shock and harvested continuously for another 14 hours. Note that PER2 oscillation seems to be coupled to that of PER1 in wt cells. Representative of two experiments. See also Figure S5.

### *Cry* genes affect circadian rhythms through posttranslational regulation of PER in addition to transcriptional regulation of *Per*

CRY proteins are considered the main transcriptional inhibitors in the circadian feedback loop (Kume et al., 1999), but circadian phase (timing of feedback inhibition) is determined by cooperation between PER and CRY in the inhibitor complex (Chen et al., 2009; Ukai-Tadenuma et al., 2011). Like the *Per* paralogs, the two *Cry* genes are not completely redundant. *Cry1* and *Cry2* KO mice show shortened and lengthened behavioral rhythms (by <1hr) compared to wt mice, respectively, suggesting that they may have a slightly antagonistic role in setting period (van der Horst et al., 1999; Vitaterna et al., 1999). To measure how they affect *Per*-reporter rhythms, *Cry1* and *Cry2* genes were singly and doubly deleted in both *Per1^Luc^* and *Per2^Luc^*KI reporter cells. Deletion of *Cry1* and *Cry2* in these cells produced period shortening and lengthening, respectively, consistent with phenotypes in mice (Figure 5A-D) (van der Horst et al., 1999; Vitaterna et al., 1999). However, the amount of alteration was more dramatic in U2OS cells compared to mice. Both PER1-Luc and PER2-Luc signals increased significantly in *Cry1* KO cells but slightly decreased in *Cry2* KO cells, suggesting that CRY1 is the stronger inhibitor than CRY2, consistent with recent studies (Fribourgh et al., 2020; Rosensweig et al., 2018). In line with the bioluminescence data, both PER1 and PER2 protein levels were elevated in *Cry1* KO cells (Figure 5E). These data elegantly explain why *Cry1* and *Cry2* KO can cause period shortening and lengthening, respectively. In *Cry1* KO cells, PER threshold levels for feedback inhibition would be reached earlier causing shortening of the feedback loop, while those would be delayed in *Cry2* KO cells resulting in period lengthening. *Cry1/2* double-KO cells did not become completely arrhythmic immediately, as opposed to immediate arrhythmicity in *Cry1/2* double-KO mice. Instead, both PER1-Luc and PER2-Luc were robustly rhythmic for one circadian cycle and then became arrhythmic (Figure 5F and 5G). The first cycle is not included because it could be zeitgeber-driven rather than an endogenous feedback loop-driven rhythm. PER1-Luc bioluminescence was higher than in wt cells, but PER2-Luc signals were slightly lower compared to wt cells. Changes in PER abundance by *Cry* KO seems to directly reflect changes in *Per* transcription in these mutant cells (Figure 5H). Consistent with above data (Figure 3 and 4), *Per1* transcription is more dramatically regulated than that of *Per2* by *Cry* KO. High and low levels of PER1 and PER2 in *Cry* double-KO cells, respectively, were confirmed by immunoblotting which also revealed an additional difference between the PER paralogs (Figure 5I and 5J). Hypophosphorylated isoforms of PER1 and PER2 were much more pronounced in *Cry* double-KO cells. Hyperphosphorylated species failed to accumulate after cycloheximide treatment (Figure S6A) suggesting that hyperphosphorylated species cannot be generated or they are very unstable without CRY. Because hyperphosphorylated PER species were readily detectable after calyculin A (CA) treatment (Figure S6B), CRY seems to protect hyperphosphorylated species from dephosphorylation or rapid degradation. Because PER, especially PER1, is more rapidly degraded in *Cry* double-KO cells after CHX treatment (Figure S6A), these data suggest that the conversion of hypophosphorylated to hyperphosphorylated PER species is normal, but they are unstable without CRY. Similar observations were made in liver tissue between wt and *Cry* double-KO mice (Lee et al., 2001). Hyperphosphorylation of PER was restored when transgenic CRY was expressed in *Cry* double-KO cells, indicating that the phenotype is not due to the genetic defect (Figure S6C).

**Fig 5.**
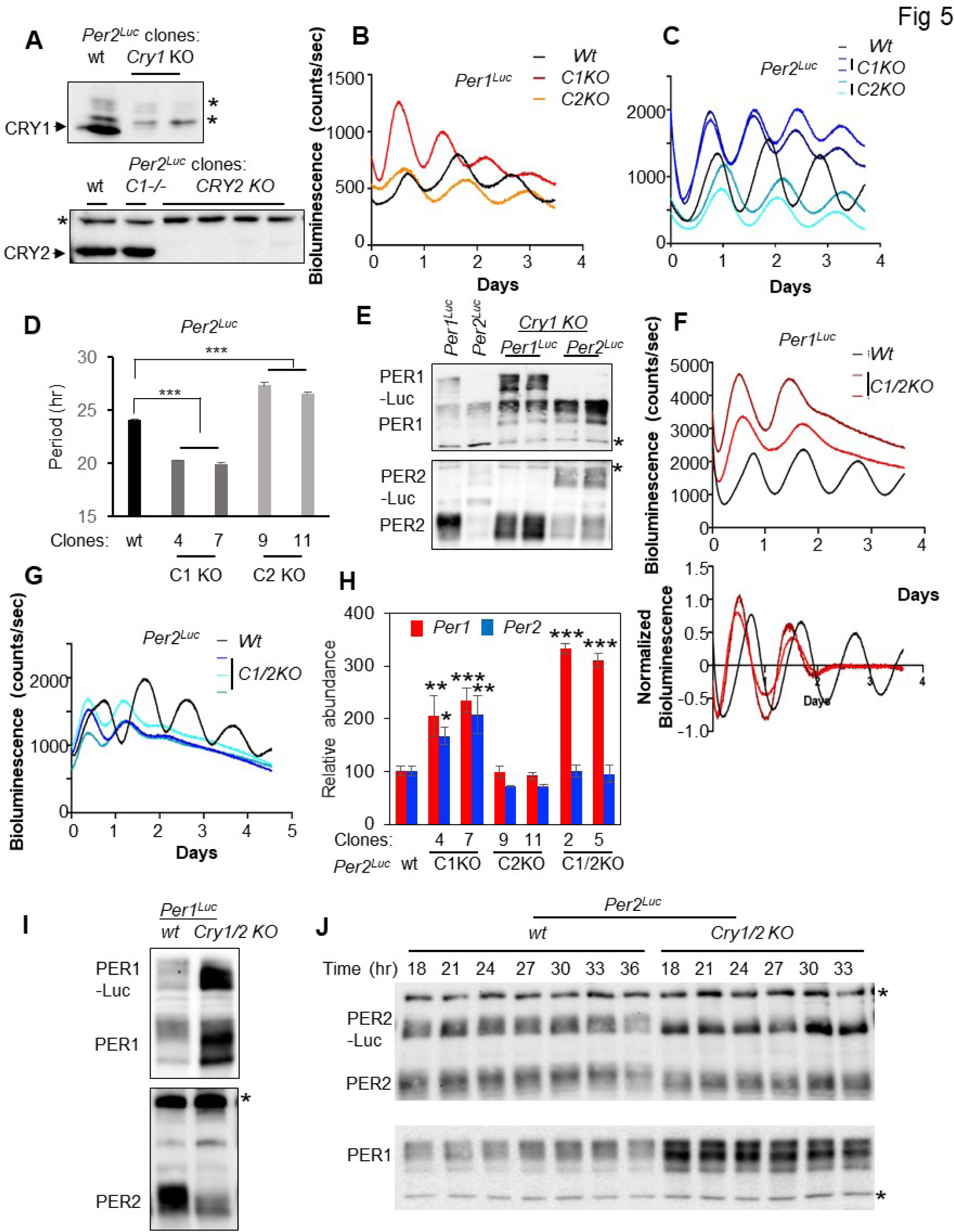
Deletion of *Cry* genes in the reporter cells reveals functional difference between two CRYs. (A) *Cry* KOs were screened by altered bioluminescence rhythms and confirmed by immunoblotting. (B-D) *Cry* KO phenotypes were consistent with mouse genetics data. Each trace represents a different clone and is representative of three traces. Mean+/-SD of two clones each for *Cry1* and *Cry2* KO in *Per2^Luc^* is shown in (D). (E) PER protein levels were elevated in *Cry1* KO cells. (F, G) PER1-Luc and PER2-Luc signals were higher and lower, respectively in *Per^Luc^* reporter cells. A normalized bioluminescence graph is shown in (F) to compare period between wt and *Cry1/2* double KO cells. (H) Modulations of PER proteins by *Cry* KO are explainable by modulations in *Per* mRNA levels. Note that *Per1* mRNA levels were elevated in *Cry1* and *Cry1/2* KO cells but those of *Per2* mRNA were significantly elevated only in *Cry1* KO cells. Mean+/-SEM of three RT-qPCR experiments. (I) Modulation of PER proteins in *Cry1/2* KO cells was confirmed by immunoblotting. (J) PER proteins are constitutively hypophosphorylated in *Cry1/2* double-KO cells. * nonspecific band. See also Figure S6.

### C-terminus of CK1δ/ε is critical for rhythm generation through making stable interaction with their substrate PER

There are seven CK1 isoforms in mammals, encoded by seven different genes (Knippschild et al., 2014). They are very well conserved in the catalytic domain (AA1 to ∼300), but highly divergent in C-terminal noncatalytic domains (Figure S7). The circadian clock is completely compromised and PER hyperphosphorylation is absent in CK1δ/ε double-KO cells (Lee et al., 2011). PER2 is not even noticeably phosphorylated in the double-KO cells based on mobility shift. Although there is also evidence that CK1δ/ε are not the only kinases that phosphorylate PER (Chiu et al., 2011; Hirota et al., 2010; Smith et al., 2008), the KO data indicate that other CK1 isoforms and other kinases cannot replace CK1δ/ε as essential regulators of PER phosphorylation for the circadian clock. Because the catalytic domains are highly conserved (Figure S7), the C-terminal regions of CK1δ/ε are likely important for their function as circadian kinases. We began our studies of CK1δ/ε in our reporter cells by confirming their circadian functionality: deleting both genes and measuring the effect on bioluminescence rhythms. In our initial attempt to generate CK1δ/ε double-KO cells, we induced frame-shift mutations in exon 4 of the *CK1δ* gene, and then targeted exon 2 in *CK1ε* in the resulting *CK1δ* KO clones (Figure S7). However, no viable colonies were detected after single-cell sorting, indicating that double KO leads to lethality, as has been reported in MEFs (Lee et al., 2011). To circumvent this issue and test functionality of the C-terminus of CK1ε at the same time, frame-shifting mutations were introduced in the non-essential C-terminal region in CK1ε (exon 8) (Figure S7). As expected, many viable colonies were obtained. Since our anti-CK1δ/ε antibodies were raised against the non-conserved C-termini, C-terminus-truncated CK1ε could not be detected on immunoblots (Figure 6A). When CK1δ or ε was deleted by targeting early exons, E4 and E2, respectively, circadian rhythms were significantly lengthened by 1.5–2 hrs (Figure 6B-D), consistent with genetic data in mice (Etchegaray et al., 2009). Interestingly, frame-shifting mutations in either exon 2 or exon 8 in the C-terminus of CK1ε produced similarly lengthened rhythms, suggesting that the C-terminus of CK1ε plays an important role in the clockwork (Figure 6C and 6D). Consistent with these data, deletion of the C-terminus of CK1ε (exon 8) in a *CK1δ* KO background (thus creating double-mutant cells) produced much longer periods than *CK1δ* single-KO cells in both *Per1^Luc^* and *Per2^Luc^*cells (Figure 6E-G and S8A), further supporting the importance of C-terminal regions in CK1δ/ε for the clockwork. One of the double mutants isolated based on period lengthening in *Per1^Luc^* CK1δ/ε double mutant clones had deletion of two AAs plus a point mutation instead of a frame-shifting mutation (*Per1^Luc^*CKKO-2) (Fig 6G bottom panel and S7). The mutant clone showed even longer period than a C-terminal deletion mutant clone *Per1^Luc^*CKKO-1 (Fig 6F, 6G and S8A). In all of these double mutant cells, PER could be hyperphosphorylated in a rhythmic manner albeit with a delay. There was accumulation of hyperphosphorylated PER species (Figure 6H), which was probably induced by their increased stability (Figure 6I). We believe the slowed hyperphosphorylation and thus increased stability are caused by attenuated interaction between PER1 and C-terminus-deleted or mutated CK1ε. PER interaction with the mutant CK1ε in the *Per1^Luc^* CKKO-2 clone was significantly attenuated compared to wt CK1ε when measured using transiently expressed proteins (Figure 6J). Overall, these data suggest that C-terminal regions of CK1δ/ε play an important role in promoting hyperphosphorylation in PER for timely degradation through making stable interactions between enzyme and substrate.

**Fig 6.**
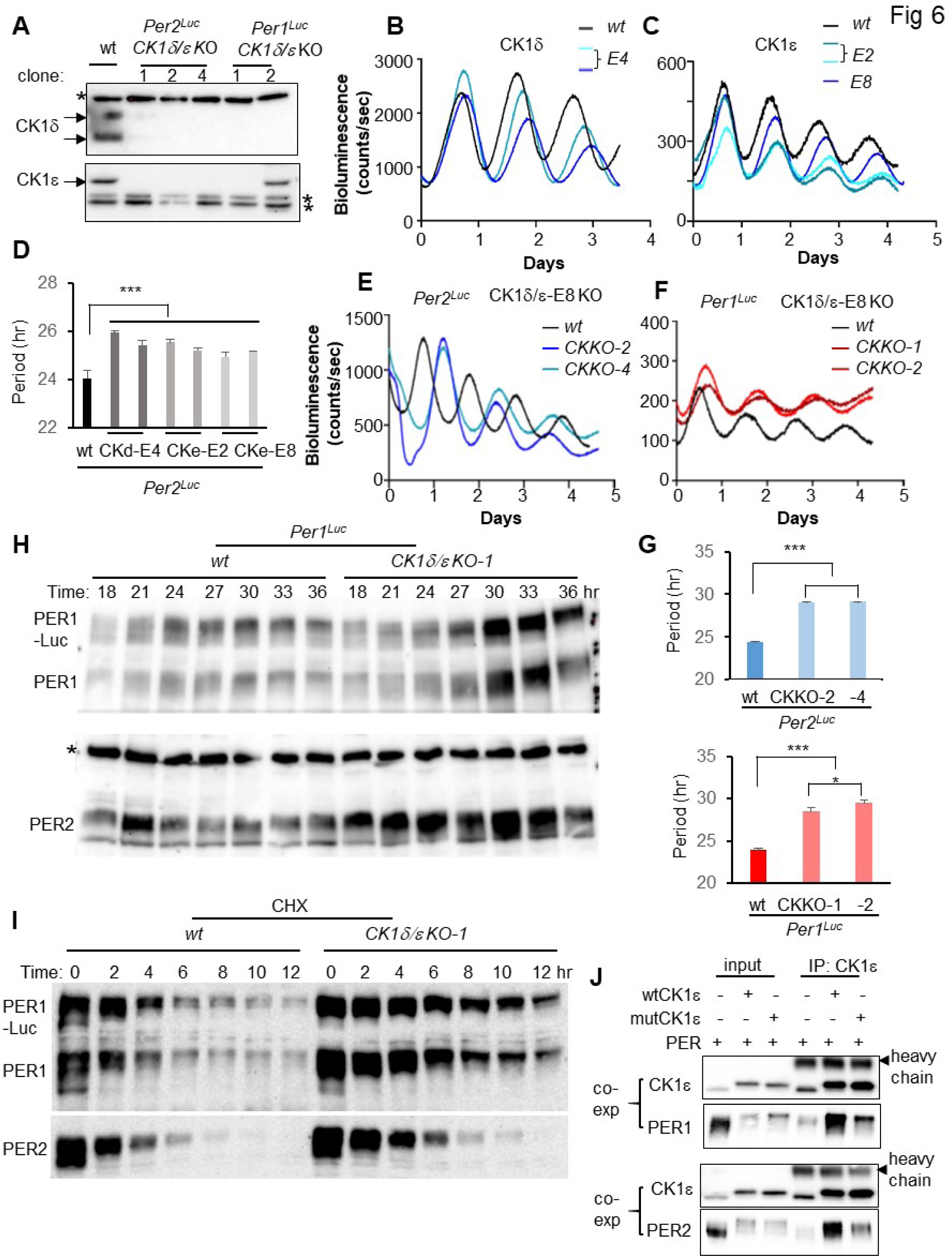
C-termini of CK1δ/ε play important roles in rhythm generation. (A) Deletion of CK1δ/ε was screened by altered bioluminescence rhythms and confirmed by immunoblotting. * nonspecific band. Note that CK1ε is detected in double KO #2 in *Per1^Luc^* reporter cell due to an in-frame mutation in one allele (*Per1^Luc^* CKKO-2) (see Figure S7). (B-D) Single KOs of *CK1δ* and *CK1ε* and C-terminus deletion of CK1ε all lead to lengthened rhythms. Two representative clones for each KO are shown in (D). Mean+/-SD, N=3. (E-G) *CK1δ/ε* double-mutant clones showed dramatically lengthened rhythms. Note that CKKO-2 in (F) represents the double-mutant clone with the in-frame mutation in C-terminus instead of a complete knockout via a frame- shifting mutation (Figure S7). Mean+/-SD, N=3. (H) Hyperphosphorylated PER species were pronounced in CK1 double mutant cells. CKKO-1 clone in *Per1^Luc^* reporter cell is shown. * nonspecific band. (I) PER, especially PER1 was stabilized in double mutant clones. CKKO-1 clone in *Per1^Luc^*reporter cell is shown. (J) PER interaction with a mutant CK1ε with two AA deletion (RE) in C-terminus was significantly attenuated. PER was co-expressed with wt CK1ε or mut CK1ε and subjected to coimmunoprecipitation (coIP). Note that small amounts of transgenic PER were copurified with endogenous CK1ε by the anti-CK1ε antibody. Transgenic CK1ε is larger than the endogenous counterpart by 18 AAs. See also Figures S7 and S8.

### The circadian phosphotimer requires stable interaction between CK1δ/ε and PER

Two main events regulated by the PER phosphotimer are nuclear entry and degradation of the PER-containing complex; these events are separated by ∼12 hours and define distinct circadian phases (D’Alessandro et al., 2017; Lee et al., 2001; Lee et al., 2011). Because PER phosphorylation by CK1δ/ε is responsible for these events directly, the timing of PER phosphorylation must occur over the same extended period, ∼12 hrs. CK1δ and ε bind the substrate PER stably through a dedicated ∼220 AA domain called CKBD in PER (Eide et al., 2005; Lee et al., 2004), which is different from a typical transient kinase-substrate relationship. Although many previous studies suggested that this domain (which, as discussed earlier, contains the FASPS mutation S662G) plays an important role in setting period, how CKBD contributes to timed phosphorylation has been unsolved. Consistent with previous studies (Narasimamurthy et al., 2018; Philpott et al., 2020), deletion of a few AAs around S662 produced shortened rhythms (Philpott, 2022). While generating small AA indels around S662 in CKBD to study the significance of the domain, a mutant clone missing 2/3 of CKBD was isolated (Figure 7A and S8B). Unlike the small AA indel mutants, this clone exhibited a significantly lengthened rhythm, ∼27.5 hrs (Figure 7A and 7B). The mutant PER2 levels were elevated compared to wt PER2. When *Per1* was deleted in this clone, rhythms were completely eliminated, and protein levels were further elevated, demonstrating that the mutant PER2 is not functional, and the mutation is semi-dominant over *Per1* (Figure 7A). The mutant PER2 was constitutively hyperphosphorylated, which was not dependent on PER1 (Figure 7C and 7D). As expected, its interaction with CK1δ/ε was dramatically attenuated (Figure S8C). Although these data indicate that PER hyperphosphorylation is not dependent on stable interaction with CK1δ/ε, when the mutant cell was treated with a specific CK1δ/ε inhibitor, PF670462, PER hyperphosphorylation was inhibited in both wt and mutant PER2, demonstrating that hyperphosphorylation of the mutant PER2 does depend on CK1δ/ε (Figure 7E and S8D). Furthermore, hyperphosphorylation of the mutant PER2 was more sensitive to the inhibitor; maximum inhibition was achieved at a much lower dose of the inhibitor compared to wt PER2 (Figure 7E and S8D). These data suggest that hyperphosphorylation of PER by CK1δ/ε does not require stable interaction, but a functional phosphotimer requires CKBD because it can allow slow and progressive phosphorylation. These data are consistent with a recent study showing that PER phosphorylation by CK1δ does not require CK1δ anchoring through CKBD (Marzoll et al., 2022). In the absence of the stable interaction, PER phosphorylation by the kinases would be done in a transient manner like typical kinase reactions. The mutant PER2 was predominantly nuclear as expected from their phosphorylation status and significantly more stable than wt PER2 (Figure 7F, 7G and S9).

**Fig 7.**
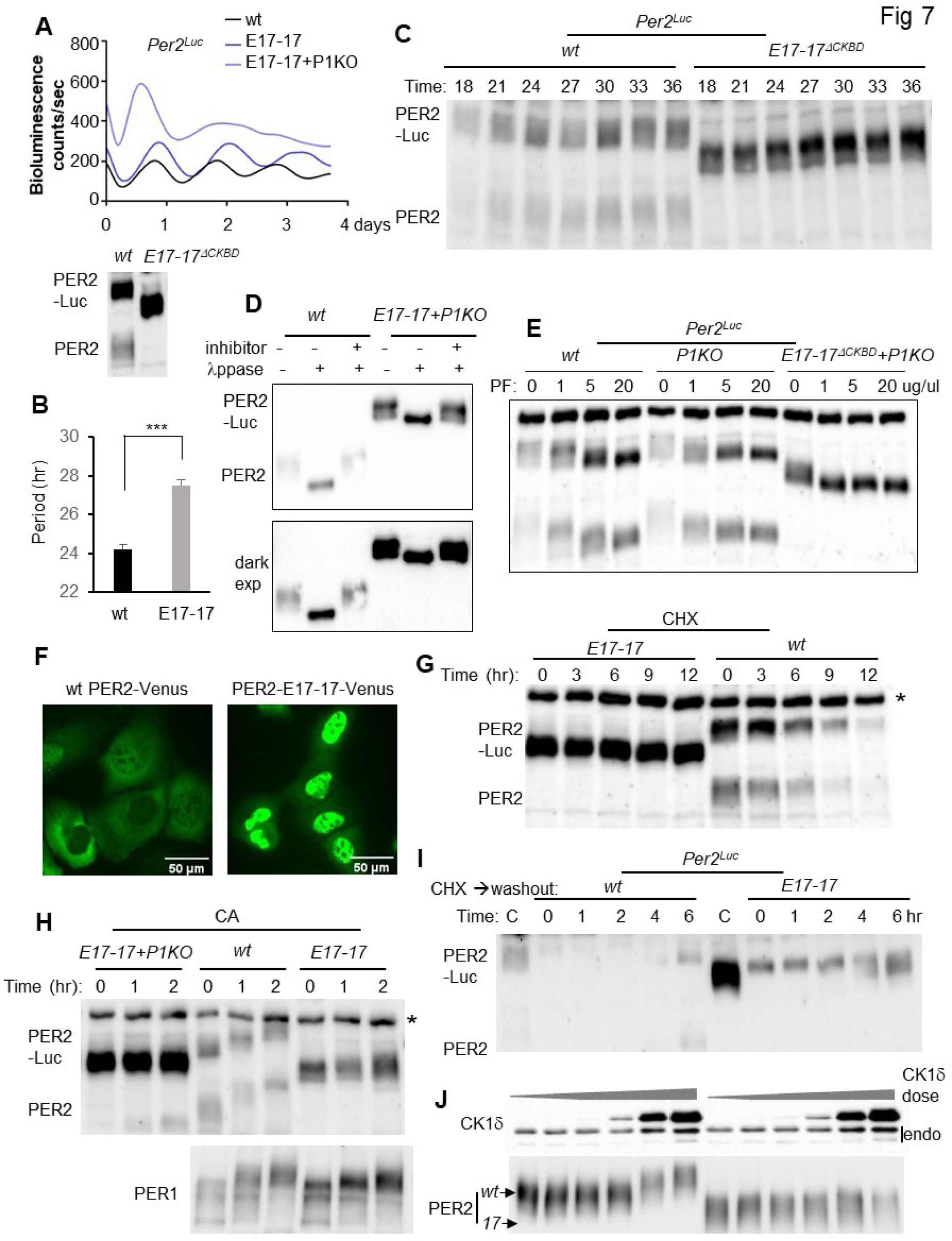
CKBD allows for slow phosphorylation kinetics essential for the phosphotimer. (A, B) A *Per2* mutant clone lacking the CKBD showed lengthened rhythms in a *Per1* wt background but arrhythmicity in *Per1* KO cells. Note that PER2-Luc signals were elevated in *Per1* wt and further elevated in the *Per1* KO background. A similar number of cells was used. A smaller size mutant protein was confirmed by sequencing of cDNA (Figure S8A) and immunoblotting (bottom panel in (A)). (C) The mutant PER2 seemed to be constitutively hyperphosphorylated based on mobility shift on the immunoblot. (D) Hyperphosphorylation was confirmed by phosphatase treatment. Two exposures are shown. Representative of two experiments. (E) Phosphorylation of the mutant PER2 is mediated by CK1δ/ε. Lower doses were used in Figure S8D. (F) The mutant PER2 was constitutively nuclear. Wt PER2 was detectable in both cytoplasm and nucleus. When wt PER2 was expressed in high amounts, they were predominantly cytoplasmic (Figure S9) (Beesley et al., 2020). (G) The mutant PER2 was significantly more stable than wt PER2. (H) The mutant PER2 was not additionally phosphorylated by CA treatment (see extra hyperphosphorylation in Fig S6B). (I) Phosphorylation of de novo mutant PER2 was significantly advanced compared to its wt counterpart. Time ‘0 hr’ represents 8 hrs after CHX treatment right before washout. Note that a significant amount of the mutant PER2 still existed after CHX treatment. (J) The mutant PER2 is less phosphorylated in vitro by CK1δ compared to wt PER2. Increasing amounts of CK1δ were co-expressed with a fixed amount of PER2. See also Figures S8 and S9.

When the mutant PER2 was treated with CA, extra hyperphosphorylated species were not detected, suggesting that this mutant PER2 is defective in this extra phosphorylation, which is normally not detected in a steady state because these extra hyperphosphorylated species are rapidly turned over by β-TRCP-regulated proteasomal degradation (Figure 7H) (D’Alessandro et al., 2017). Faster phosphorylation kinetics for the mutant PER2 was confirmed with de novo PER2 after existing PERs was depleted (Figure 7I). The lack of extra hyperphosphorylation was reproduced with transiently expressed CK1δ and mutant PER2 (Figure 7J).

## Discussion

A critical bottleneck in studying complex biological systems like the circadian clock is the lack of an efficient in vivo-like platform where endogenous genes can be easily manipulated to test diverse hypotheses in a time- and cost-effective manner. Historically, manipulation of endogenous clock genes in cell culture models required first developing mutant mouse models from which cells such as MEFs were then harvested (D’Alessandro et al., 2017; Park et al., 2015; Xu et al., 2015). However, recent developments in CRISPR genome editing have created new opportunities for generating cell culture models without first generating mutant mice. Several studies including ours demonstrated that clock genes can be knocked out efficiently in culture using CRISPR (Jin et al., 2019; Kim et al., 2021; Korge et al., 2015; Lu et al., 2016; Matsu-Ura et al., 2015). One key limitation in cell models today is the lack of precise endogenous phase and period reporters working like clock hands other than the *mPer2-Luc* reporter in MEFs.

Fluorescence reporters have been developed (Gabriel et al., 2021), but they are far less accurate than bioluminescence reporters and require heavy deconvolution of data, especially when signal is barely above background. When we put mRuby3 before luciferase to produce the PER- mRuby3 fusion protein, fluorescence signal was significantly lower than non-fusion mRuby3 and barely above background when measured by FACS and microscopy. This is probably due to decreased stability of mRuby3 when it is attached to the unstable PER. We were able to observe robust rhythms for more than 10 days from our bioluminescence reporter cells compared to 2-3 days reported using fluorescence reporters (Gabriel et al., 2021).

Fusion of a fairly large protein, luciferase, to PER did not seem to affect phosphorylation and stability of PER, which suggests a dominant regulation of PER at the posttranslational level and can explain a wt-like circadian phenotype in the *mPer2^Luc^* KI mouse. Although PER1 and PER2 are independently rhythmic in abundance and phosphorylation and redundant in generation of circadian rhythms, their kinetics such as stability and speed of phosphorylation differ significantly. Because PER protein phase and thus phase in the feedback loop would be determined by these properties, it would be expected that phase of clock-controlled genes would be different between *Per1* and *Per2* KO cells. Indeed, the phase of *Bmal1-Luc* was reversed in *Per1* KO, but not in *Per2* KO cells, suggesting that *Per1* phase is dominant over that of *Per2* in wt cells, probably due to higher abundance and leading phosphorylation kinetics. However, the circadian phase of the master clock in the SCN is not altered in *Per1* KO mice. Both *Per1* and *Per2* KO mice show similar phases in behavioral rhythms in constant darkness after LD entrainment (Pendergast et al., 2010). We believe the difference between peripheral and master clocks is due to different entrainment mechanisms. In the master clock, both *Per* genes are rapidly induced by photic signals resulting in acute phase shifting (Reppert and Weaver, 2001). However, in peripheral cells, *Per2* rhythm is not rapidly reset by zeitgebers because its transcription is not acutely induced while *Per1* rhythm is acutely reset by these signals through rapid induction of mRNA as in SCN by photic signals (Balsalobre et al., 2000). This has a significant implication in humans with a defective *Per1* gene because their peripheral clocks could be dissociated from the SCN clock.

Because PER-Luc oscillates with a circadian period in *Cry1/2* KO cells at least for one complete cycle (Jin et al., 2019; Putker et al., 2021), PER alone seems to act as a feedback inhibitor, but the clock cannot be sustained without CRY. PER can directly interact with CLOCK:BMAL1 and repress the activity of the complex in in vitro reporter assays supporting the inhibitory role (Chen et al., 2009; Kume et al., 1999). In *β-Trcp* mutant cells, the clock cannot be sustained because hyperphosphorylated PER species are too stable in the nucleus (D’Alessandro et al., 2017). Similarly, it may be that the clock cannot be sustained in *Cry1/2* KO cells because hyperphosphorylated PER species are too unstable, and CRY is necessary to extend nuclear presence of PER and PER-containing inhibitory complexes.

Our data suggest that the progressive and controlled phosphorylation of PER is mediated by stable interaction between the kinase and substrate resulting in slowing down kinase movement (processivity) on PER for serial phosphorylation. The phosphotimer can be also modulated by mutations in CK1δ/ε. CK1s are considered anion- or phosphate-binding kinases because basic amino acids (AAs) in the anion-binding pockets interact with anions or phosphate groups on substrates (Flotow et al., 1990; Venerando et al., 2014). CK1s depend on negative charges or prior phosphorylation on a substrate for processive phosphorylation as in a typical consensus motif pSxxS (Flotow et al., 1990; Philpott et al., 2020; Venerando et al., 2014), which appears throughout the entire PER1 and PER2 protein, not just the FASPS domain, yet current working models of the clockwork focus on only two motifs, FASPS and a degron motif (Narasimamurthy and Virshup, 2021; Philpott et al., 2020). CK1s have several substrate-facing, anion-binding pockets; thus the more PER is phosphorylated, the more robustly PER will hold CK1, resulting in slower processivity. In vivo, CK1δ/ε are predominantly copurified only with hyperphosphorylated PER species (Lee et al., 2001), supporting the model. Further, charge inversion mutations such as *tau* (R178C) and K224D in one of the anion-binding pockets of CK1, or mutations or small deletions in the CK1-binding domain in PER generally accelerate the phosphotimer and the clock (Narasimamurthy and Virshup, 2021; Philpott, 2022; Shinohara et al., 2017). Consistent with this model, deletion of the whole CKBD resulted in typical rapid kinase reactions, leading to constitutive hyperphosphorylation. This level of phosphorylation was enough for nuclear entry, but not for degradation as shown in Fig 7. The PER phosphotimer seems to have two distinctive phases, one for nuclear entry and the other for degradation, matching with onset and offset sleep phases. CKBD is required for the second phase of the phosphotimer. It is not clear whether CKBD itself contains a critical phospho-degron or whether stable CK1 binding to PER is required for phosphorylation in a degron somewhere else in PER.

The phosphotimer is at the heart of the circadian clock; it is the basis for how a cell can measure time precisely over timeframes much longer than typical cellular processes. Our studies revealed many critical properties of this timer using diverse clock mutants. In addition, our human reporter cell model could provide direct insights into pathogenicity of many period- and phase-altering mutations in CK1 and PER and serve as an efficient platform to test diverse hypotheses developed via human genetics and biochemical studies.

## Acknowledgements

We thank Dennis Chang for assistance with manuscript revisions and Robert Tomko for gel filtration chromatography. We thank Jonathan M. Philpott and Carrie L. Partch for critical reading of the manuscript and providing feedback. This work was supported by NIH R01 GM131283 awarded to C.L. and by Korea Institute for Advancement of Technology (KIAT, P0002007, HRD Program for Industrial Innovation) grant to H.S.

## Author contributions

Conceptualization: C.L., H.S. and J.P.; Investigation: all authors; Formal analysis: C.L., J.P and K.L.; Writing-Original fradft: C.L., J.P. and K.L.; Writing-Review and Editing: C.L., J.P., K.L. and H.S.; Visualization: C.L., J.P. and K.L.; Supervision: C.L. and H.S.; Funding Acquisition: C.L. and H.S.

## STAR Methods

### CONTACT FOR REAGENT AND RESOURCE SHARING

*Further information and requests for resources and reagents should be directed to and will be fulfilled by the Lead Contact, Choogon Lee (*).

### EXPERIMENTAL MODEL AND SUBJECT DETAILS

#### Cell lines

The U2OS cell line was purchased from ATCC (#HTB-96).

U2OS-Bmal1-Luc wt and *Per2* knockout (KO) (E5-2) lines were described previously (Jin et al., 2019). U2OS-Bmal1-Luc; *Per1* KO clones were selected from the previous study (Jin et al., 2019), and #E6-2-4 was used for this study.

*mPer2^Luc^* knockin (KI) reporter MEFs were described previously (Yoo et al., 2004). HEK293a (ThermoFisher #70507) was used in Fig 6J.

All cells were maintained at 37 °C, 5% CO2 in DMEM, supplemented with 10% FBS. <u>Generation of *Per* KI cell lines</u>or all CRISPR-induced mutations, sgRNAs were selected via CHOPCHOP (https://chopchop.cbu.uib.no), cloned into pAdTrack-Cas9-DEST, and tested for efficiency by T7E1 assay as described previously (Jin et al., 2019).

Wild-type (wt) U2OS cells were transfected with all-in-one pAdTrack-Cas9-DEST plasmids (sgRNA sequence described in Supplementary T'
able 1 and T7E1 primers described in Supplementary table 2) and linear repair templates (Supplementary Fig 10 and 11) using jetOPTIMUS according to the manufacturer’s protocol (jetOPTIMUS transfection reagent, Polyplus). The linear templates were prepared from Per-Luc-T2A-Ruby_pUC19 plasmid (Supplementary Fig 12) by PCR amplification. Cloning primers for the plasmids are described in Supplementary Table 3 and Supplementary Figure 12. Assembly of these amplicons were done using NEB HiFi DNA Assembly Master Mix according to the manufacturer’s protocol.

The transfected cells were maintained and expanded for 10 days before they were subjected to trypsinization and FACS sorting using BD FACSAria SORP equipped with an Automated Cell Deposition Unit (ACDU) for mRuby3-positive single-cell sorting into 96 well plates as well as bulk sorting. The single cells were expanded for 2 weeks and split into one well in a regular 24-well plate and one well in a black-wall 24-well plate (PerkinElmer #1450-605, Arkon, OH, USA). The bulk sorted cells were also added into the black-wall plate before it was set up into Lumicycle 96. Several clones with robust circadian bioluminescence rhythms were selected for each *Per* gene and subjected to junction PCR, immunoblotting with anti-PER1 (GP62), anti-PER2 (hP2-GP49) and anti-Luciferase (Luc) antibodies, and Sanger sequencing.

For Luc immunoblotting, novel polyclonal anti-Luc antibodies were generated by Cocalico Biologicals, Inc (449 Stevens Rd, Reamstown, PA) in guinea pigs using Luc AA 300-550 peptide. The antisera were validated against transfected Luc and Luc KI cells. GP77 was used in this study. To verify a single insertion of the dual reporter, frameshift mutations were introduced in early exons in these clones using all-in-one CRISPR adenovirus as described previously (Jin et al) (Fig S3). Disappearance of mRuby3 signal was confirmed by fluorescence microscopy and by FACS and bioluminescence in the Lumicycle 96. Final positive clones are summarized in Supplementary Table 1.

Mutations of clock genes in *Per1^Luc^*and *Per2^Luc^* reporter cells

Heterozygous KI clones, H10 for *Per1^Luc^*and LH1 for *Per2^Luc^*, were used to generate mutations in other clock genes. The mRuby3-expressing reporter cells were transfected with all-in-one pAdTrack-Cas9-DEST plasmids expressing GFP. In Fig S3, the reporter cells were infected with all-in-one CRISPR adenovirus to induce indels in the majority of the cells with near 100% efficiency as described previously. For clonal isolation of clock mutant cells, GFP-positive cells were sorted by FACS into 96-well plates, and these clones were further selected based on alterations in period and/or phase in bioluminescence rhythms. The sgRNA sequence and final clones are summarized in Supplementary Table 1. In each project, the majority of putative mutant clones showed a similar degree of period lengthening or shortening in bioluminescence screening. Some of these mutant clones were fully characterized by immunoblotting and sequencing.

#### *Per* mutant clones in *Per^Luc^*reporter cell lines

All-in-one adenovirus expressing CAS9 and *Per1*-E6 sgRNA used to generate indels in the *Per1* gene were described previously (Jin et al., 2019). For Fig 4C, single *Per1* KO clones were isolated by FACS followed by immunoblotting and a clone #*Per1*-E6-11 was used. To generate all-in-one adenovirus targeting *Per2* exon 6 or exon 17, the following sgRNA sequence was cloned into the pAdTrack-Cas9-Dest and adenoviruses were packaged as described previously (Jin et al., 2019).

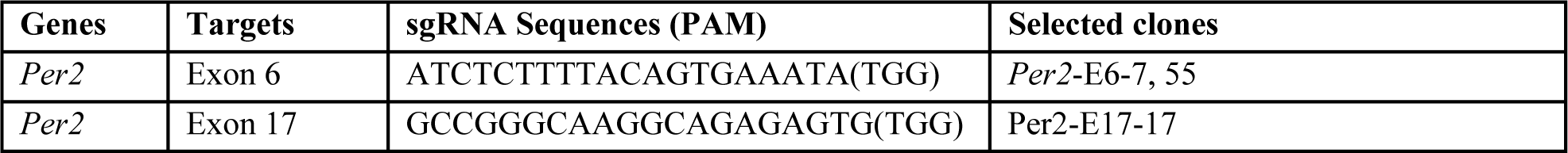

For Fig 4B, single *Per2* KO clones were isolated by using *Adv-Per2-E6* as described above. Clone #*Per2*-E6-55 was used. To generate in-frame indels in *Per2* exon 17, the *Adv-Per2-E17* was infected into the *Per2^Luc^* reporter cells and mRuby3-positive cells were sorted into 96-well plates by FACS. These cells are either wt or in-frame mutant cells because mRuby3 will be eliminated by out of frame mutations. One of the mutants was E17-17 which is missing 2/3 of CKBD. To delete *Per1* in this clone, *Adv-Per1-E6* was infected into the cells and *Per1* KO was confirmed by immunoblotting.

### METHOD DETAILS

#### Genotyping

For *Per* KI clones, KI and non-KI alleles from each clone were PCR-amplified using the primers described in Supplementary Table 4 and Supplementary Fig 13, and the amplicons were sequenced using primers described in Supplementary Table 5.

For mutant clones with indels, amplicons of the target regions were prepared using the surveyor primers (Supplementary Table 2) and sequenced. In the majority of the clones, two different Sanger sequencing traces were mixed due to different indels in two alleles. These results were deconvoluted by a computer algorithm called DECORD v3 (https://decodr.org/) into two separate traces. Accuracy of the deconvolution was confirmed by TA cloning of several amplicons from several mutant clones. Frameshift mutations in one or both alleles were also confirmed by immunoblotting.

#### Bioluminescence recording

Cells were plated into 24-well plates or 35 mm dishes to be approximately 90% confluent 24 hours prior to the start of the experiment. Immediately before the start of the experiment, cells were given a two-hour serum shock with 50% horse serum in DMEM or 10μM forskolin in DMEM (Fig S5), washed with phosphate-buffered saline (PBS) and fresh DMEM supplemented with 1% FBS, 7.5mM sodium bicarbonate, 10 mM HEPES, 25 U/ml penicillin and 25 μg/ml streptomycin. Also 0.1mM luciferin was added. The plates were sealed with cellophane tape and the dishes with cover glass with vacuum grease, and placed into a Lumicycle 96 or 32 (Actimetrics, Wilmette, IL). For all bioluminescence experiments, the results were reproduced in at least two independent experiments. Real-time levels, period, and phase of the bioluminescence rhythms were evaluated using the Lumicycle software (Actimetrics).

#### Drug treatments in cells

Cells were seeded in 60-mm dishes for immunoblots or 24-well plates for bioluminescence monitoring to be 90% confluent 24 hours prior to the experiment. For cycloheximide treatment, 8 μg/ml was added to cells and cells were collected at specified times after the treatment or placed into Lumicycle 96 for bioluminescence monitoring. For CHX washout experiments, cells were treated with CHX for 8 hrs followed by the normal DMEM, and cells were harvested at the indicated times after the CHX treatment. PF670462 was purchased from Sigma. 20 nM Calyculin A (CA, EMD chemicals) were used for cells.

#### Immunoblotting, gel filtration and Immunoprecipitation

The cells in 6 cm dishes were harvested and flash-frozen on dry ice. Protein extraction and immunoblotting were performed as previously described (D’Alessandro et al., 2015). Briefly, cells were homogenized at 4°C in 70 ml extraction buffer (EB) (0.4M NaCl, 20mM HEPES (pH 7.5), 1mM EDTA, 5mM NaF, 1 mM dithiothreitol, 0.3% Triton X-100, 5% glycerol, 0.25mM phenylmethylsulfonyl fluoride, 10mg of aprotinin per ml, 5mg of leupeptin per ml, 1mg of pepstatin A per ml). Homogenates were cleared by centrifugation 12 min, 12,000g at 4°C. Supernatants were mixed with 2x sample buffer and boiled. Proteins were separated by electrophoresis through SDS polyacrylamide gels and then transferred to nitrocellulose membranes. Membranes were blocked with 5% non-fat dry milk in TBS-0.05% Tween-20 (TBST), incubated with primary antibodies overnight followed by incubation with secondary antibodies for 1 h. The blots were developed using an enhanced chemiluminescence substrate (WestFemto, ThermoFisher Scientific).

Antibodies to clock proteins were generated as previously reported (Lee et al., 2001; Lee et al., 2004). CLK-1-GP, BM1-2-GP, C1-GP (CRY1), C2-GP (CRY2), CK1δ-GP and CK1ε-GP antibodies were used at 1:1,000 dilution in 5% milk–TBST solution. Rabbit anti-ACTIN antibody (Sigma, A5060) was used at 1:2,000.

Cell extracts from wt U2OS and a heterozygous *Per1^Luc^* reporter cells were fractionated by Superose 6 column (Cytiva, Marlborough, MA) and immunoblotted for PER1 and PER2 as described previously (Beesley et al., 2020). Fractions after the void volume of the column are shown.

Immunoprecipitation was performed as described previously (Lee et al., 2001). Briefly, protein extracts from cells harvested from 10 cm dishes were prepared as described above. 10% of the initial protein extract was saved for the input. 20μL Protein G Sepharose 4 Fast Flow beads (GE Healthcare) per reaction was equilibrated with 500μL of EB for 15 minutes on a rotating wheel. The beads were centrifuged at 3000 rpm for 15 seconds and the supernatant was removed. This wash step was repeated three additional times. After the final wash, two volumes of EB were added to the beads. To pre-clear the extracts, 10μL of the equilibrated bead solution were added to the extracts and incubated for 30 minutes on a rotating wheel at 4°C. The samples were centrifuged at 12,000 rpm for 5 minutes at 4°C and then the pre-cleared extract was transferred to a fresh tube. Then 0.1μg of affinity-purified antibody and 10 ml of the equilibrated beads were added to the tube. This mixture was incubated at 4°C on a rotating wheel for four hours. The tubes were centrifuged at 3000 rpm for fifteen seconds, and the supernatant was removed. 1mL of EB was added to the tube and incubated on a rotating wheel at 4°C for twenty minutes. The samples were centrifuged at 3000 rpm for 15 seconds, and the supernatant was removed. This step was repeated three more times to completely wash the beads. After the final wash, the majority of the EB was removed and 30μL of 1X sample buffer was added. The samples were boiled at 95°C for 3 minutes.

#### Quantitative RT–PCR

Total RNAs were extracted from U2OS wt and mutant cells using RNeasy Mini Kit (QIAGEN, Germantown, MD). After cDNAs were synthesized from 1 μg of total RNAs using qScript cDNA SuperMix (Quantabio, Beverly, MA 01915), real-time PCR was performed using SYBR Green according to the manufacturer’s protocol (PowerTrack SYBR Green Master Mix, Thermo Fisher Scientific) and primers described in the table below. RNA expression levels were calculated from three experiments.

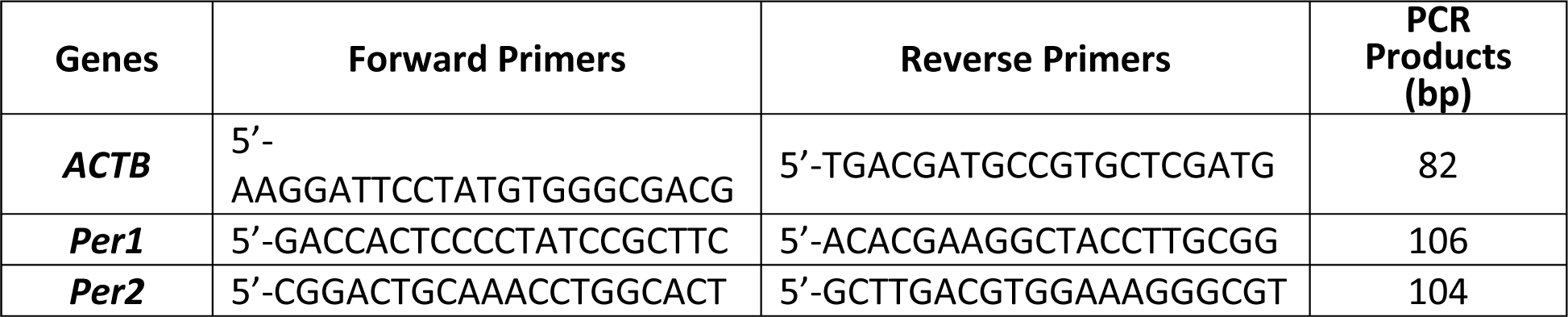

#### Adenoviral vectors and plasmids

To compare subcellular localization between wt PER2 and PER2^ΔCKBD^ (Per2-E17-17) (Fig 7F), inducible adenoviruses expressing wt hPER2-Venus or hPER2^ΔCKBD^-Venus were generated by replacing the mouse *Per2* cDNA in the previous inducible tetO-*Per2*; CMV-rtTA-pAdTrack plasmid (Beesley et al., 2020) with wt *hPer2* or *hPer2* cDNA missing the same CKBD sequence: pAdTrack-iPer2-Venus and pAdTrack-iPer2^ΔCKBD^-Venus. Adenoviruses were packaged as described previously (Jin et al., 2019).

For Fig 6J, human CK1ε cDNA was cloned into pUC19 plasmid along with CMV promoter, SV40 polyA sequence, and GFP. The T2A cleavage sequence was inserted between CK1ε and GFP resulting in CMV-CK1ε-T2A-GFP-SV40 polyA in pUC19 or pUC19_CK1ε-T2A-GFP. Expression and cleavage of the fusion protein will generate CK1ε (Fig S7) with additional 18 AAs at the C-terminus (LGGRGSLLTCGDVEENPG). The C-terminal mutant CK1ε was generated by deleting sequence encoding two missing AA (RE) in the CK1ε mutant clone. *pcDNA-Per1*, *Per2* and *CK1δ* plasmids were described previously (Chen et al., 2009).

For Fig 7J, equal amounts of pAdTrack-iPer2-Venus and pAdTrack-iPer2^ΔCKBD^-Venus plasmids were transfected into wt U2OS cells. On the following day, different titers of adenovirus expressing CK1δ were infected into the transfected cells; 0, 0.005, 0.05, 0.5, 5 and 50 MOI. Adenovirus expressing wt CK1δ was generated by cloning cDNA encoding mouse CK1δ into pAdTrack plasmid and packaging in 293A cells as described previously (Jin et al., 2019).

### QUANTIFICATION AND STATISTICAL ANALYSIS

#### Statistics

In this study, asterisks indicate significant p-values as follows: *, p<0.05; **, p<0.01; ***, p<0.001. Data across multiple experiments are shown as mean SD in all graphs except Fig 5H where mean+/-SEM is shown. One-way ANOVA and Dunnett’s post-hoc tests were used in Fig 5H. Student’s t-test was used to compare between wt and mutant cells. One-way ANOVA and Dunnett’s post-hoc tests were used for Fig 2C.

---

**Supplementary Table 1.**
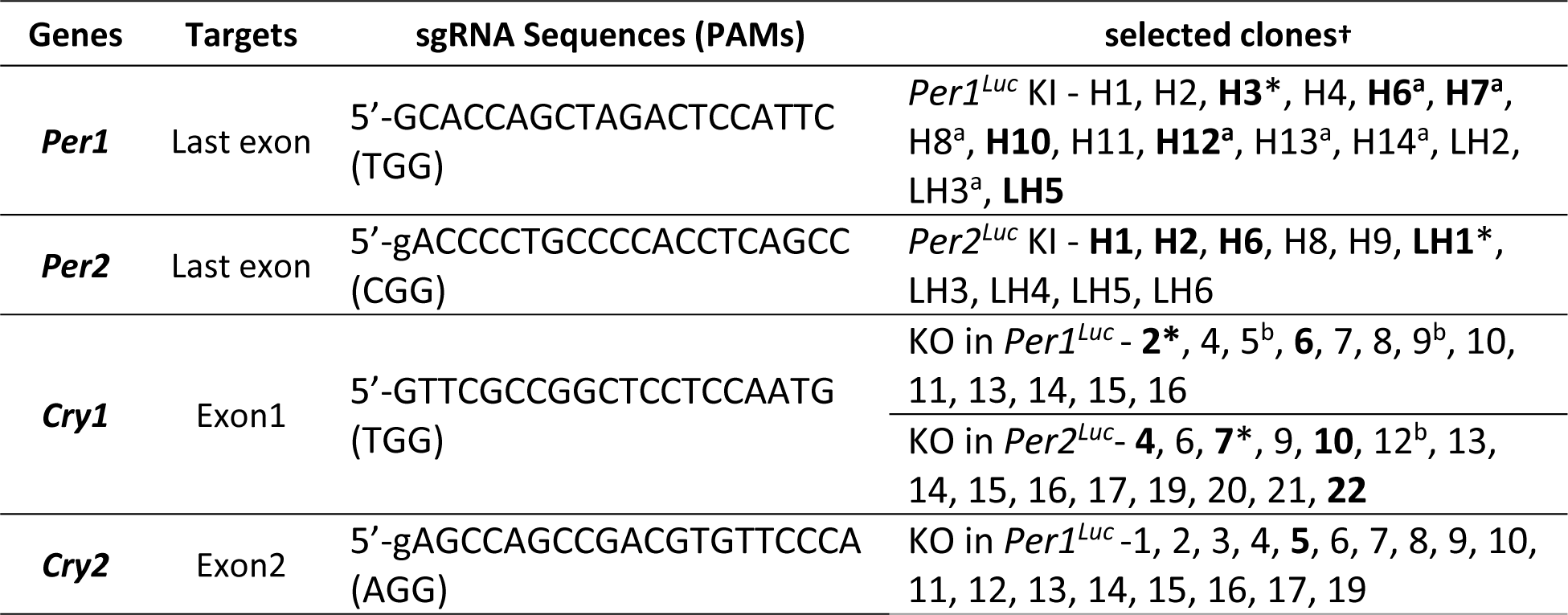
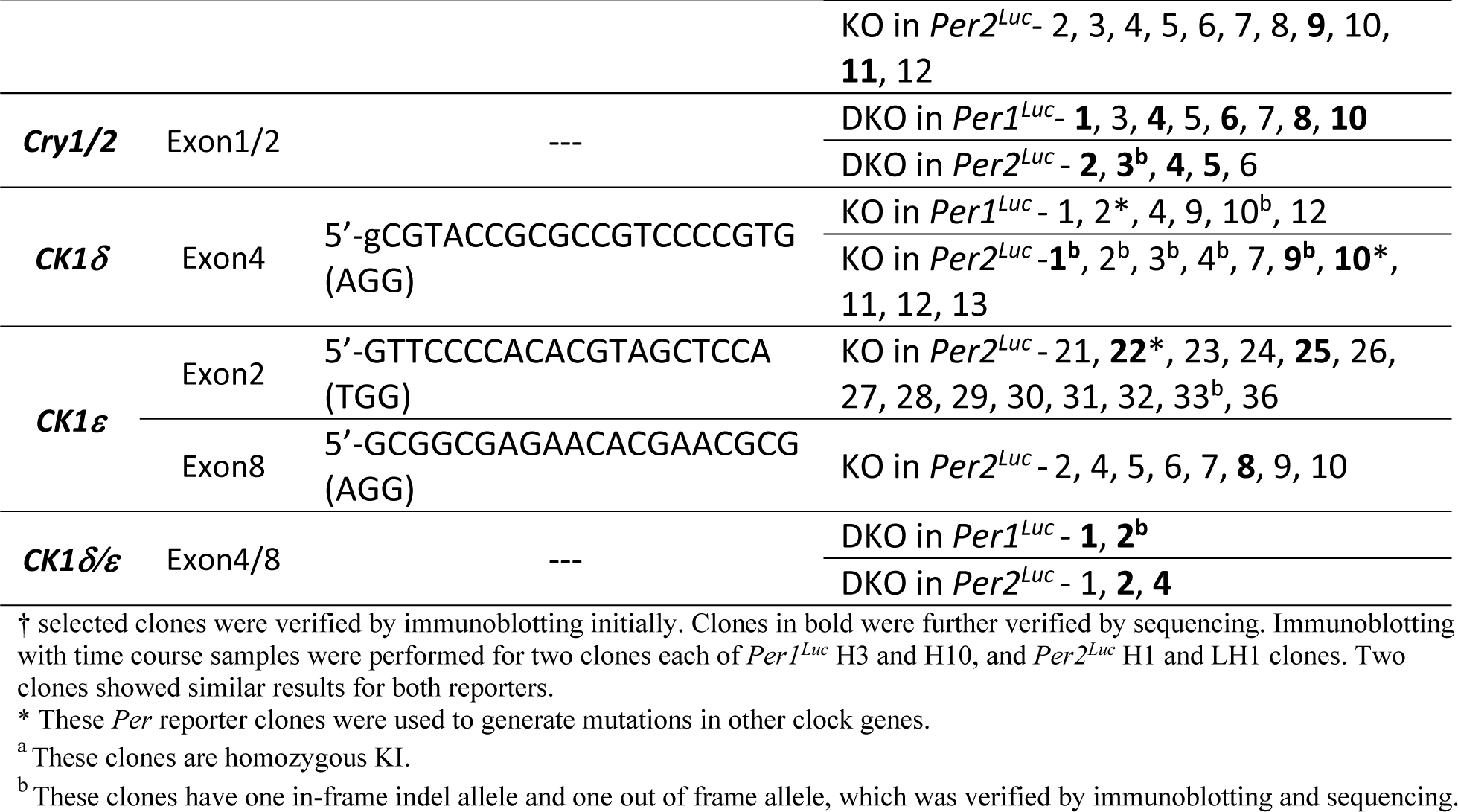
List of sgRNA Sequence and selected clones

**Supplementary Table 2.**
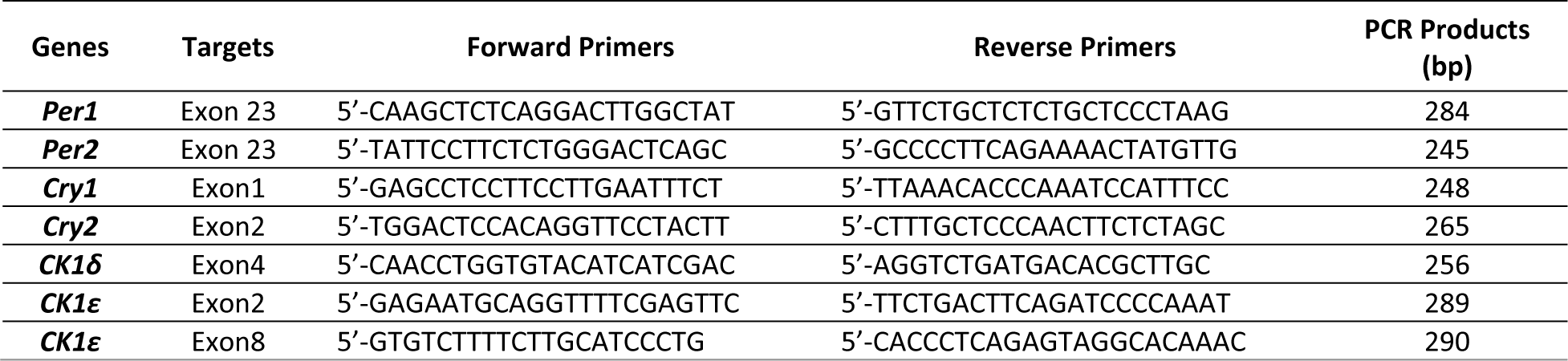
List of T7E1 Assay Primers

**Supplementary Table 3.**
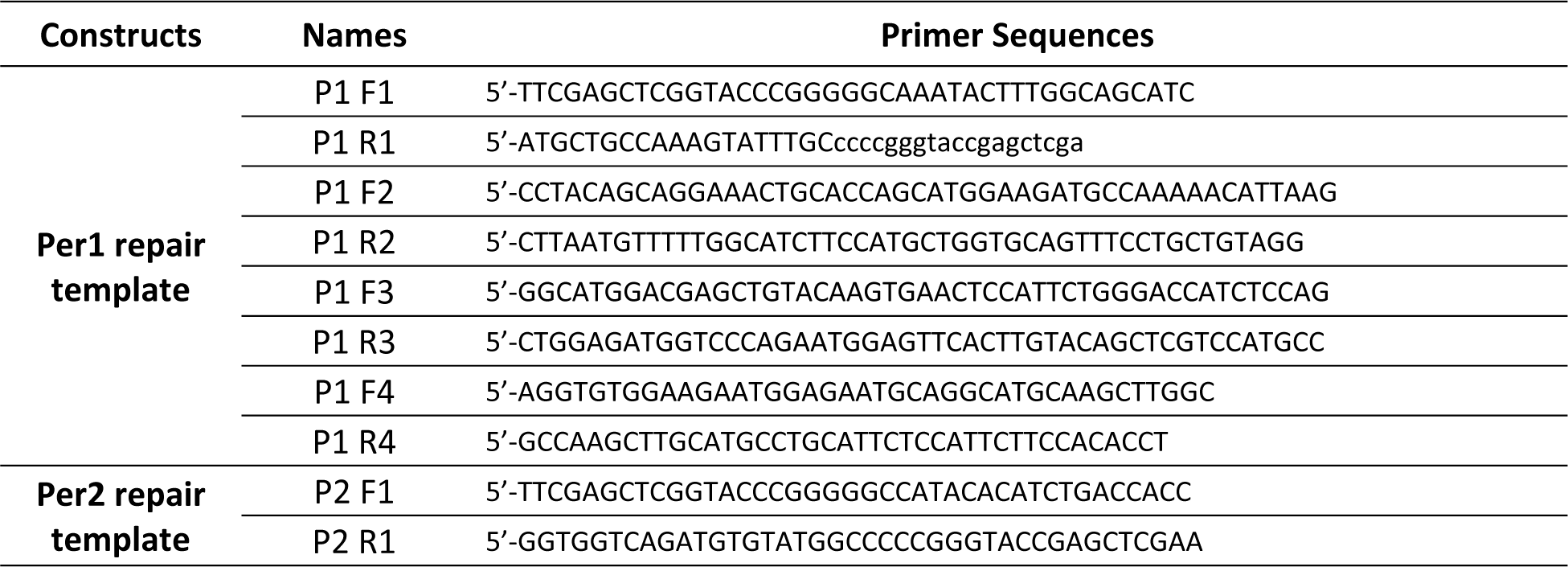
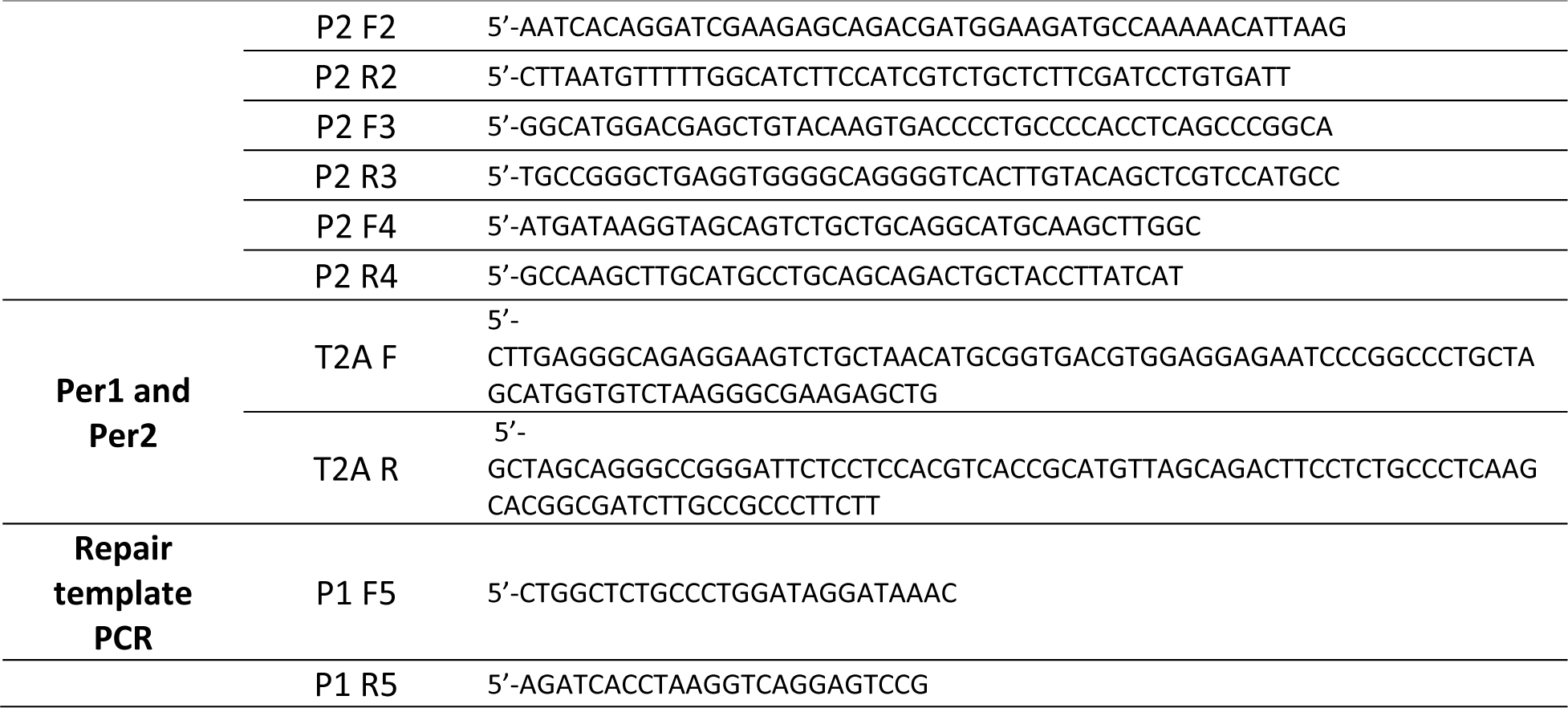
List of cloning primers for *Per* Repair Templates

**Supplementary Table 4.**
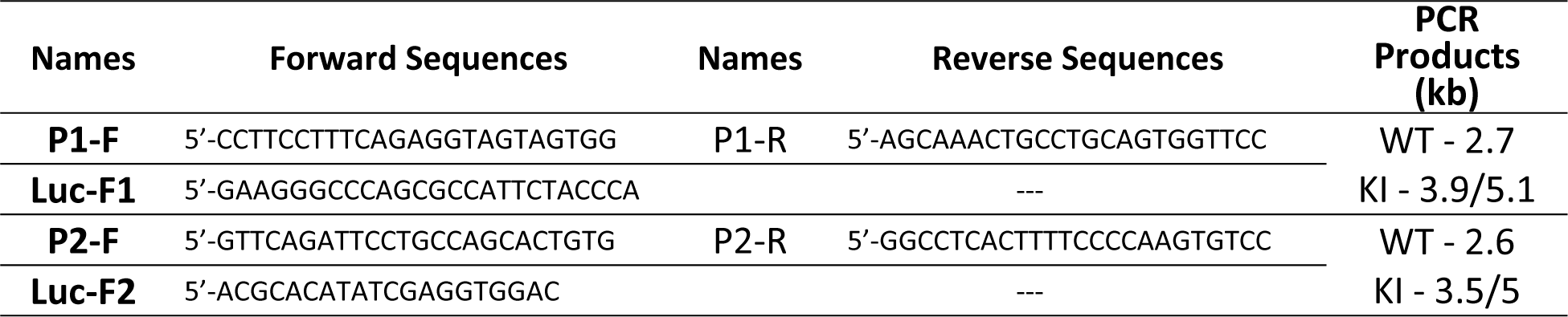
List of PCR primers for junction PCR in Fig.1D

**Supplementary Table 5.**
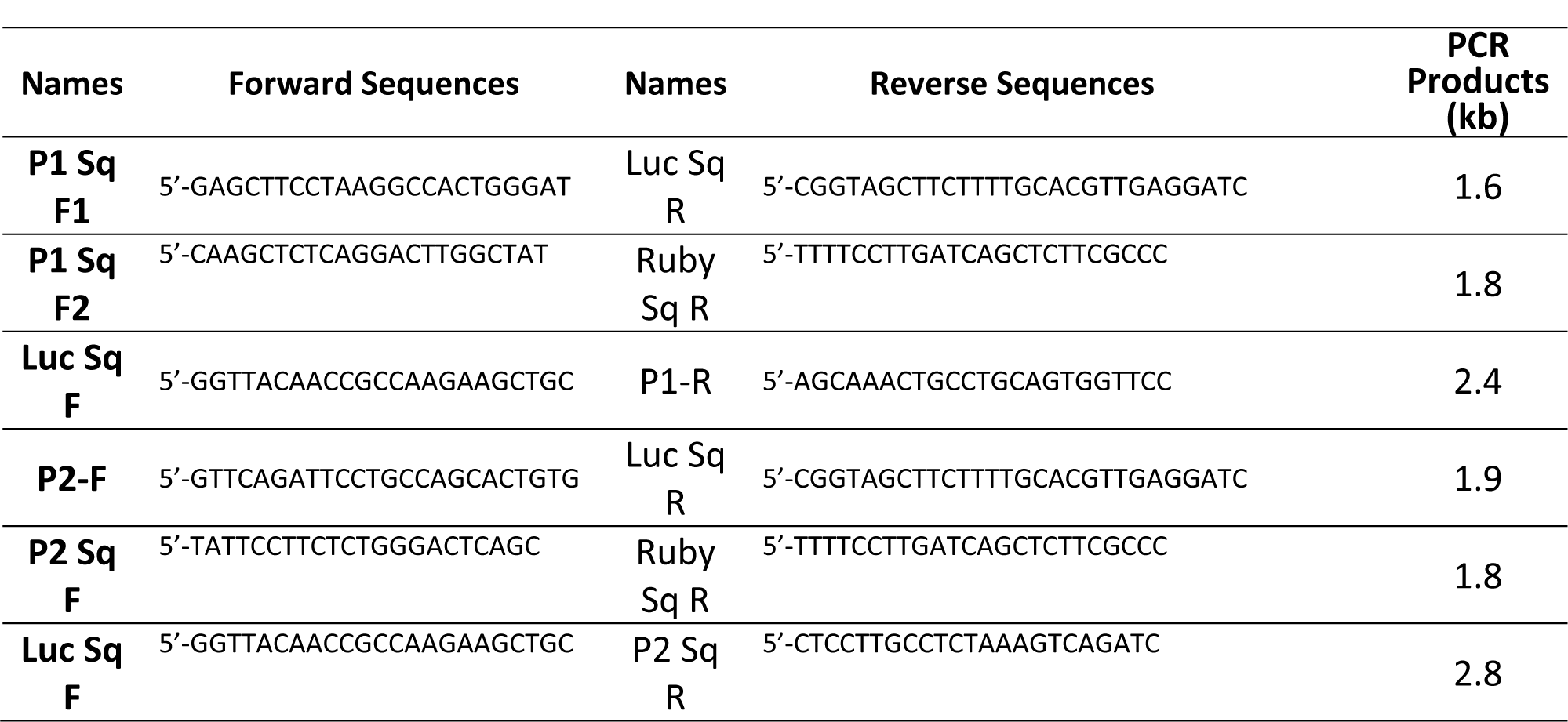
List of sequencing primers for amplicons in Supplementary Table 4.

**Supplementary Fig 10.**
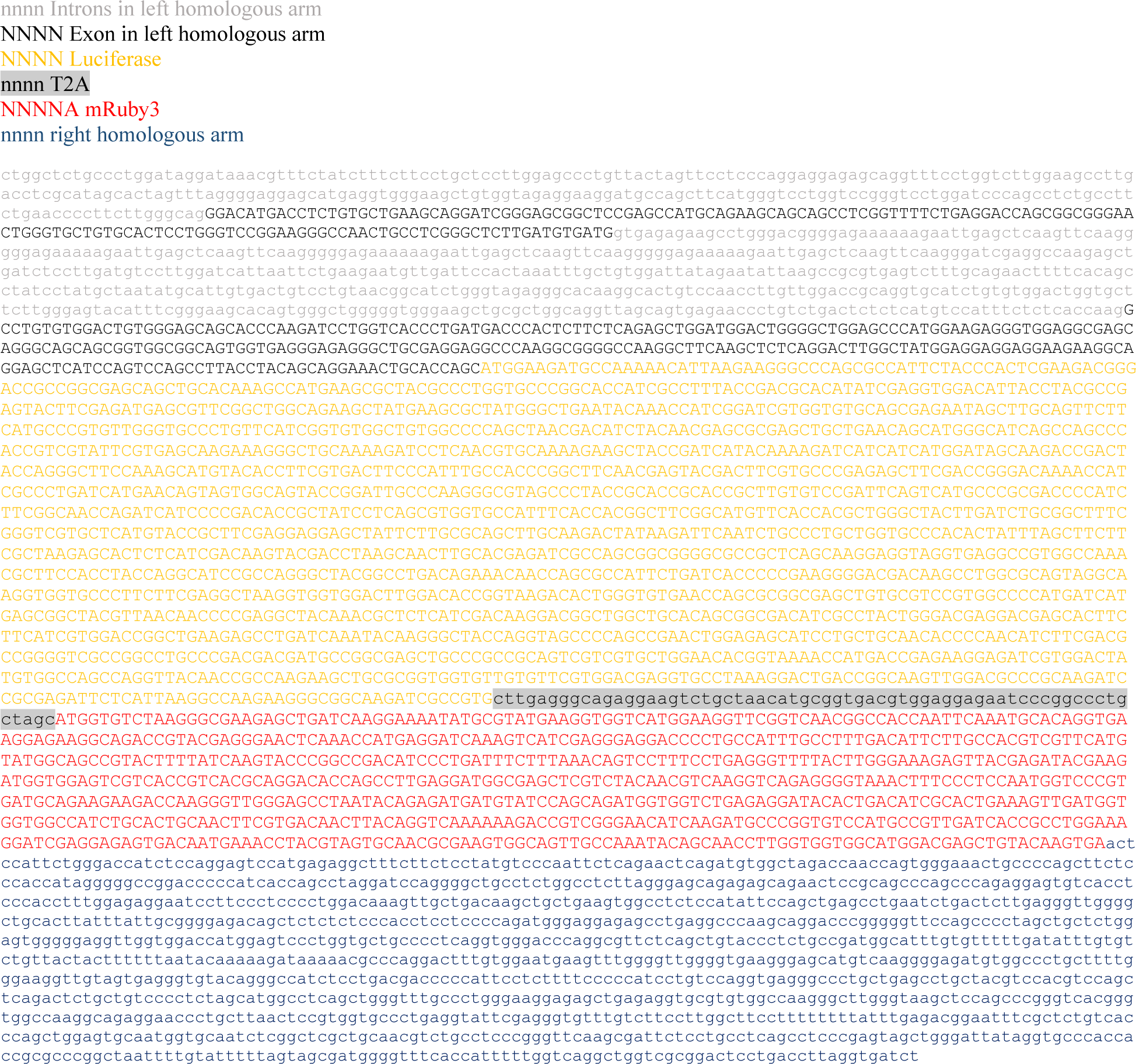
Full repair template sequence for Per1-Luc-T2A-mRuby3

**Supplementary Fig 11.**
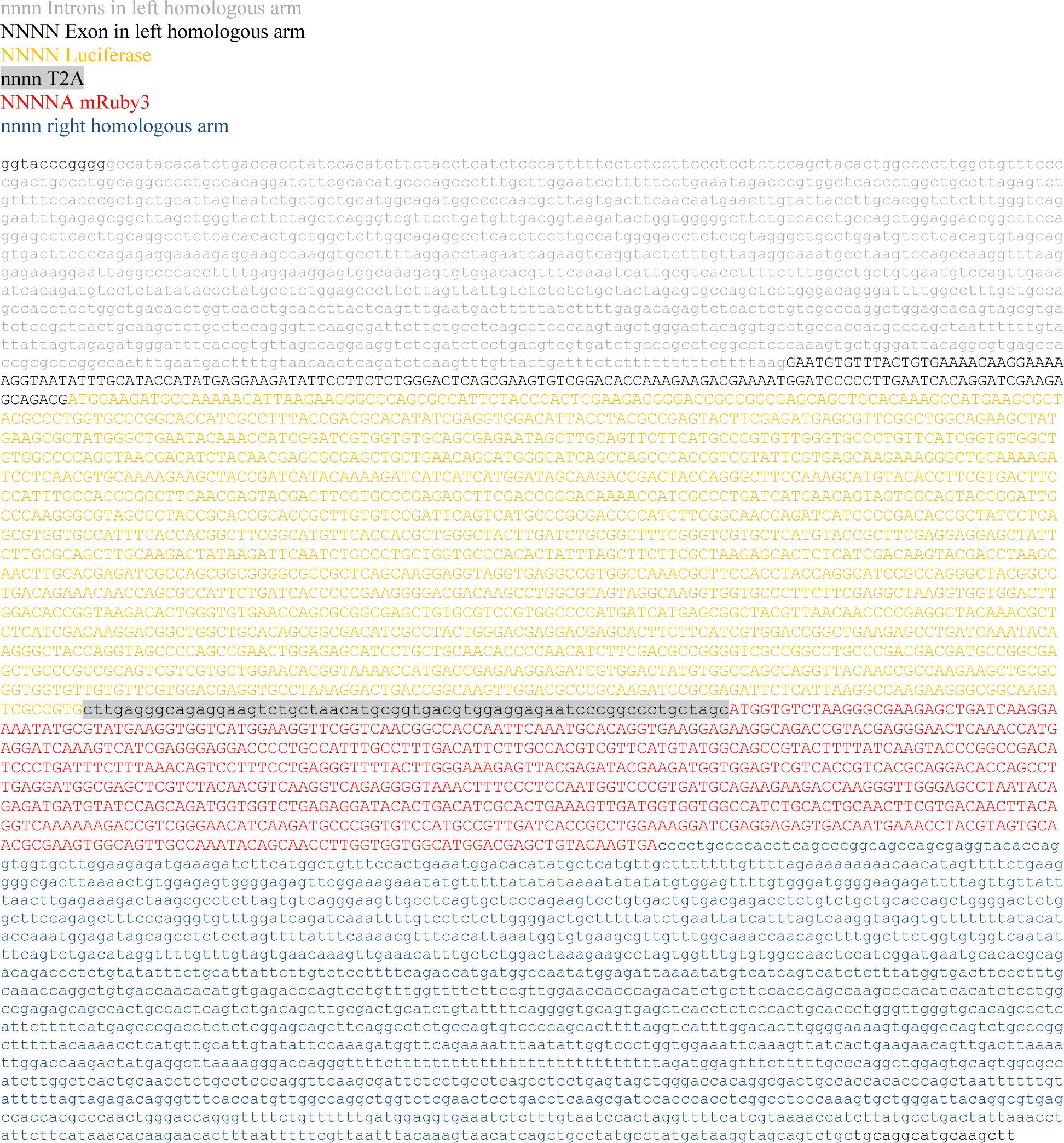
Full repair template sequence for Per2-Luc-T2A-mRuby3

**Supplementary Figure 12.**
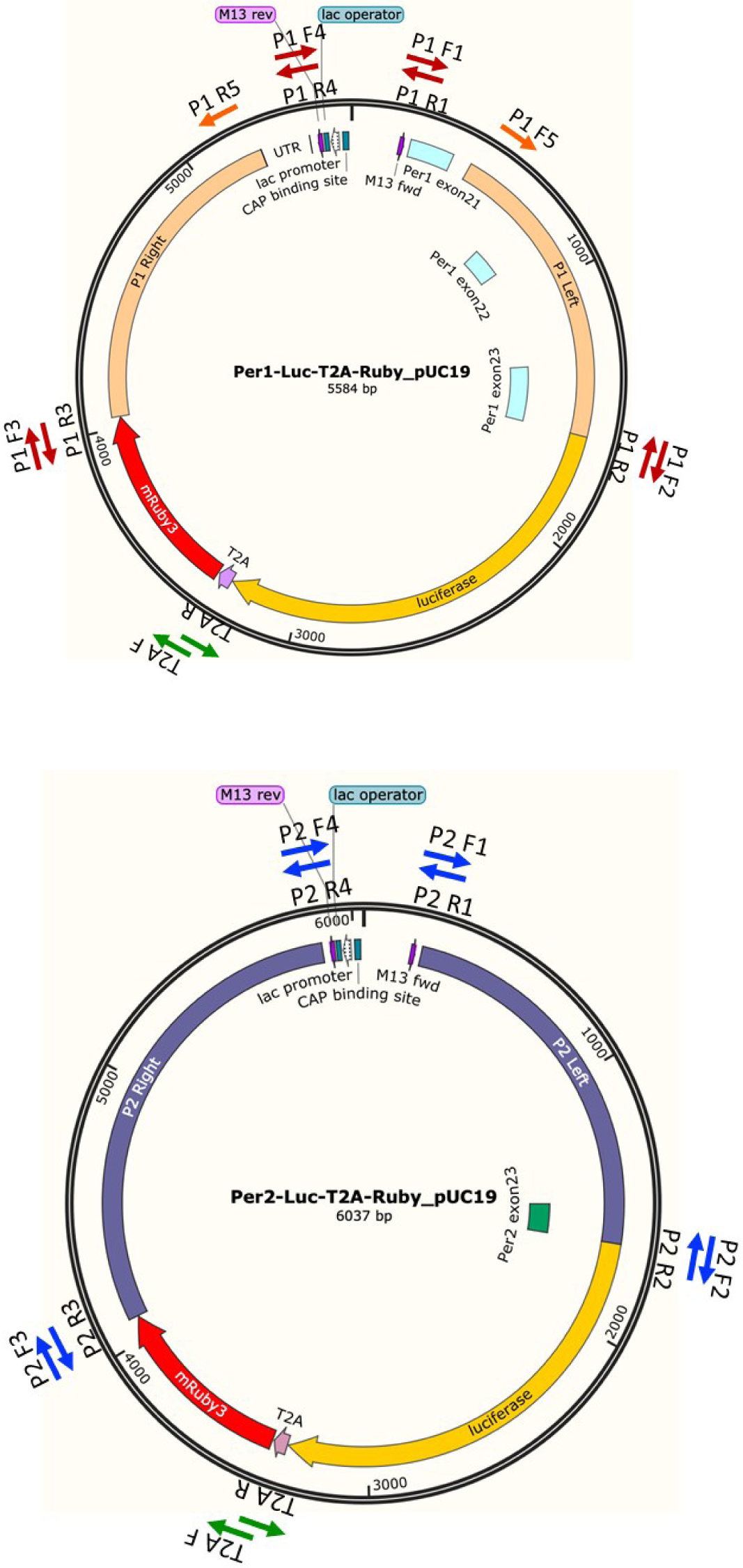
Cloning strategy for *Per* repair templates.

**Supplementary Figure 13.**
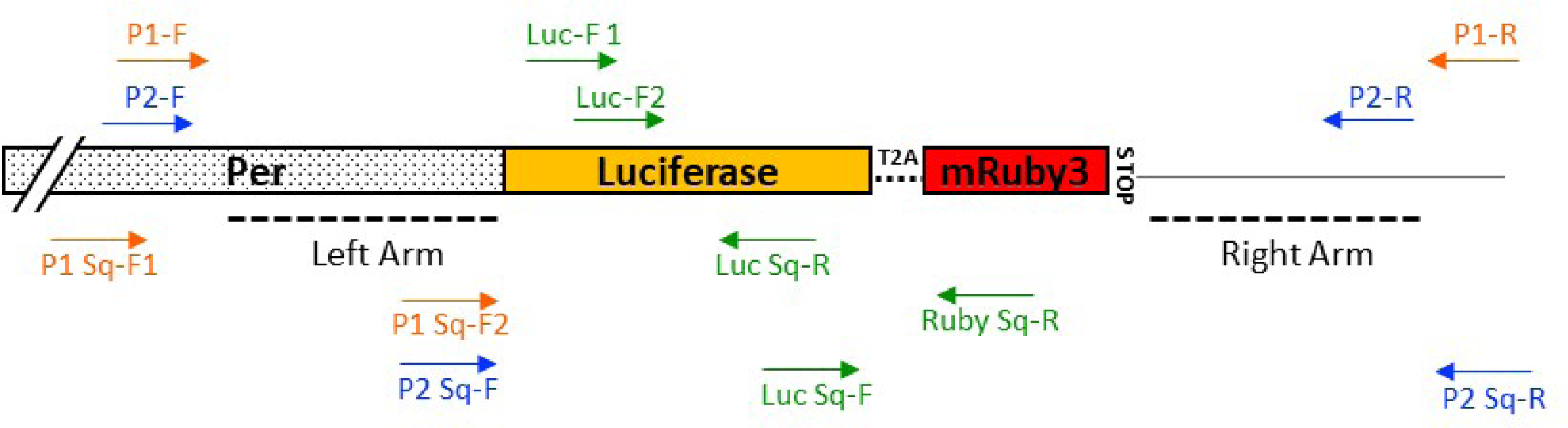
Location of primers used in Supplementary table 4 and 5.

Supplementary Information contains 9 Supplementary figures.

