## Supplementary data for "Endogenous circadian reporter cell lines as an efficient platform for studying circadian mechanisms"

Fig S1

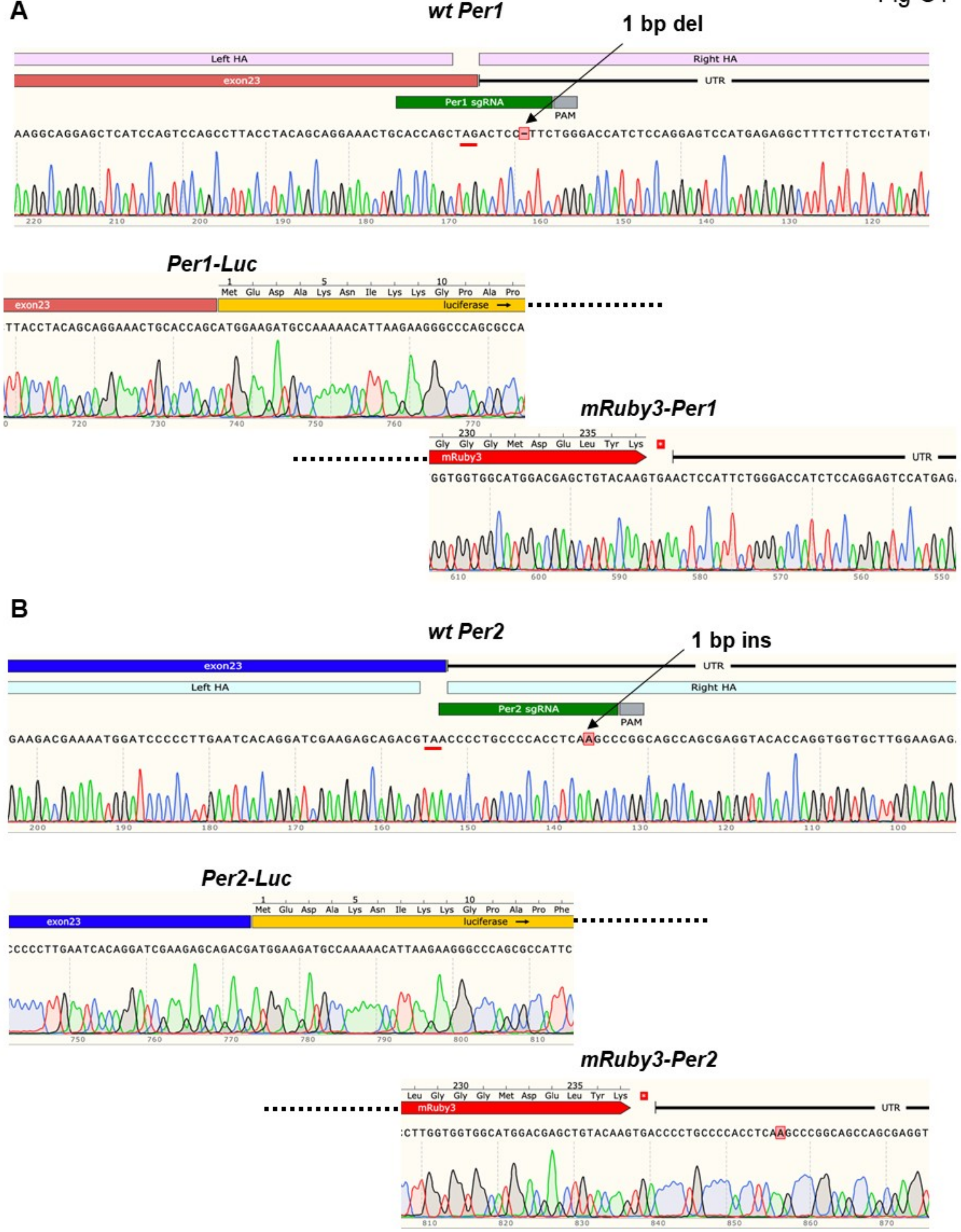


**Figure S1. Sanger sequencing of the KI alleles confirms on-target KI.** (A, B) One each of *Per1* (H3) and *Per2* (LH1) heterozygous KI clones used throughout the studies are shown. Stop codons are underlined with red. Note that a base pair (bp) was deleted in the non-KI allele in the *Per1* clone and 1 bp insertion in the *Per2* clone. There are no mutations in the KI alleles. Related to Figure 1.

Fig S2

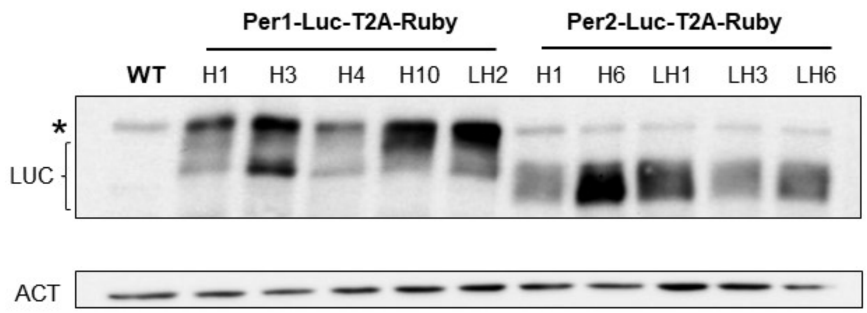


**Figure S2. Immunoblotting with anti-Luc antibody confirms specific KI.** * nonspecific band. Related to Figure 1.

Fig S3

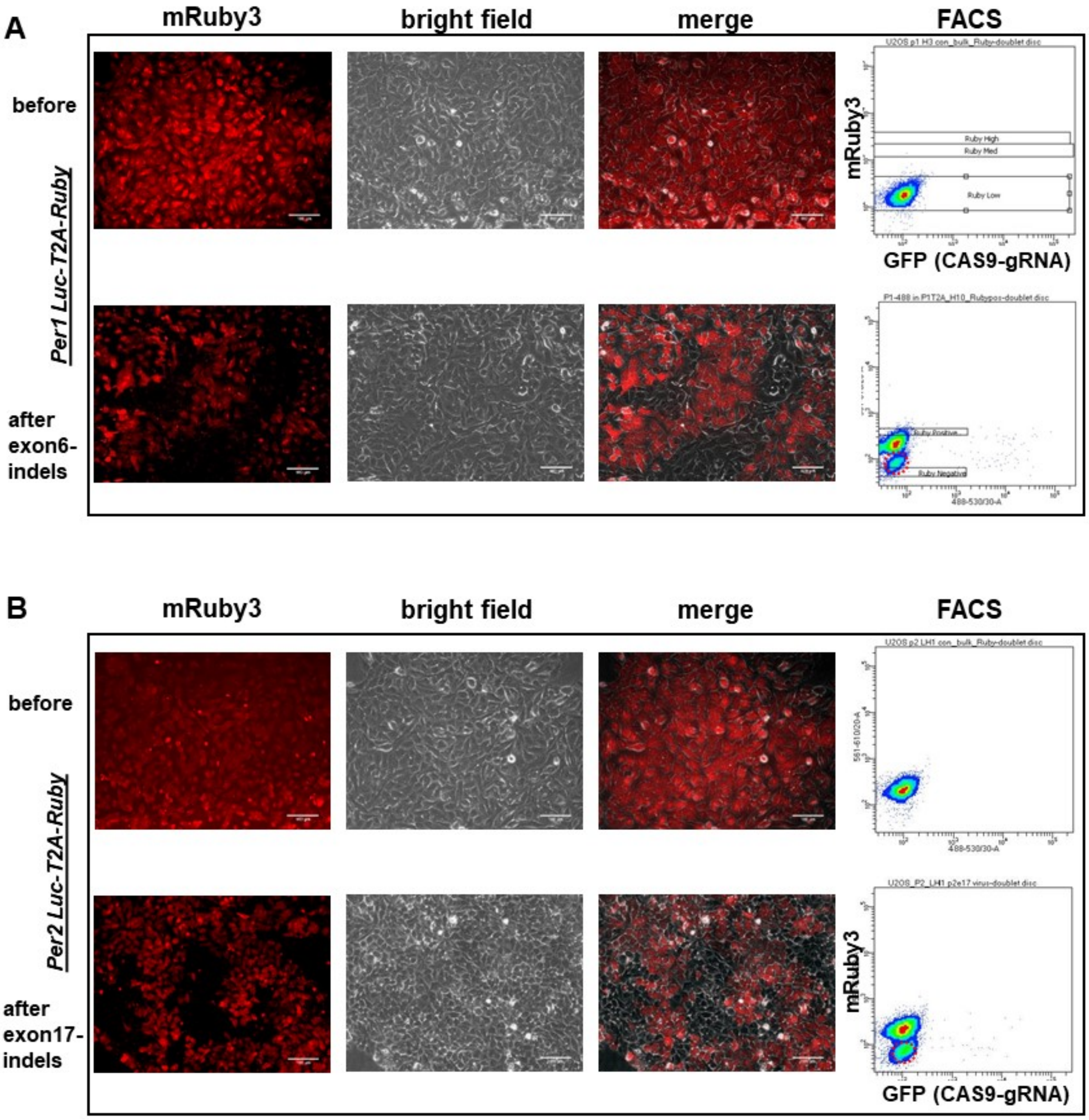


**Figure S3. mRuby3 and Luc signals disappear due to indels in early exons in *Per-Luc* reporter clones.** Representative *Per1* and *Per2* clones are shown. Indels were induced using all-in-one adenoviruses targeting exon 17 in *Per2* and exon 6 in *Per1*. To ensure 100% infection in U2OS cells with the all-in-one adenoviruses, the cells were infected at MOI of 50 twice, two days apart.

Note that a group of mRuby3-negative cells appeared after the treatment in the bottom FACS graph. These negative cells also lost bioluminescence signal. Related to Figure 1.

Fig S4

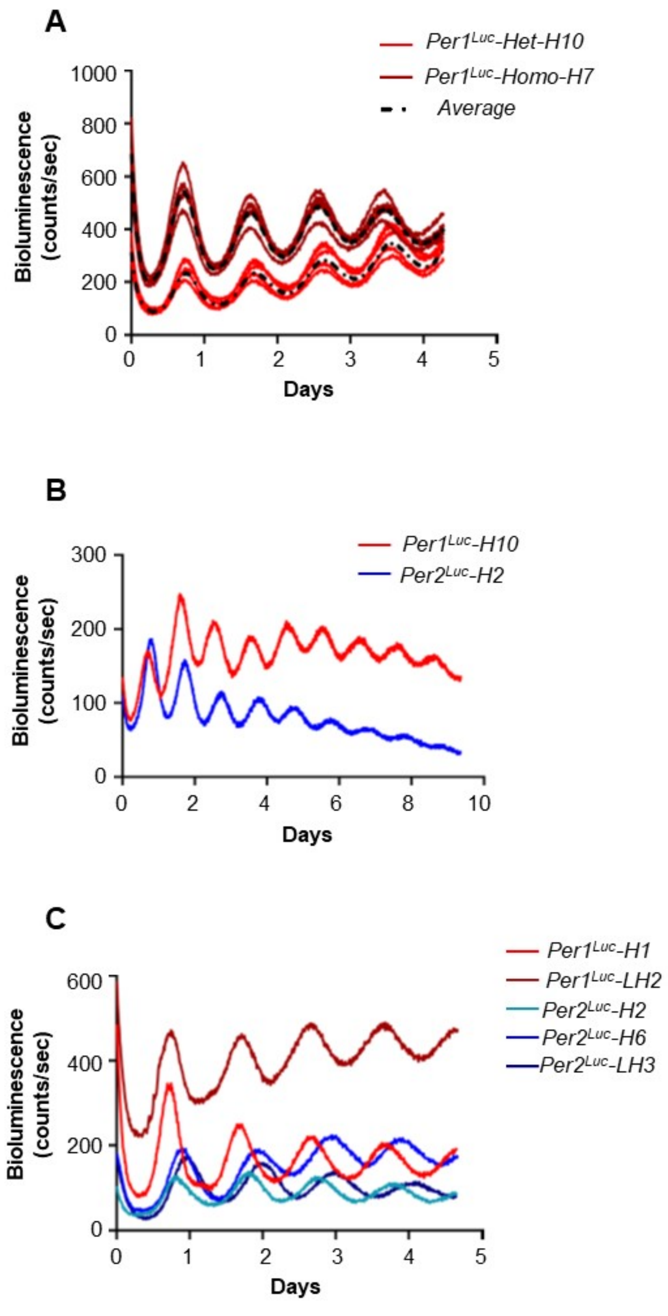


**Figure S4. Bioluminescence circadian reporters are highly quantitative and robust.** (A) A heterozygous *Per1*$^{Luc}$ was compared with a homozygous *Per1*$^{Luc}$ clone. (B) Bioluminescence rhythms were robust over 10 days. Representative *Per1*$^{Luc}$ and *Per2*$^{Luc}$ reporters are shown. (C) Circadian period was not significantly different between *Per1*$^{Luc}$ and *Per2*$^{Luc}$ reporter clones. Representative of three traces per clone is shown.
Related to Figure 2.

Fig S5

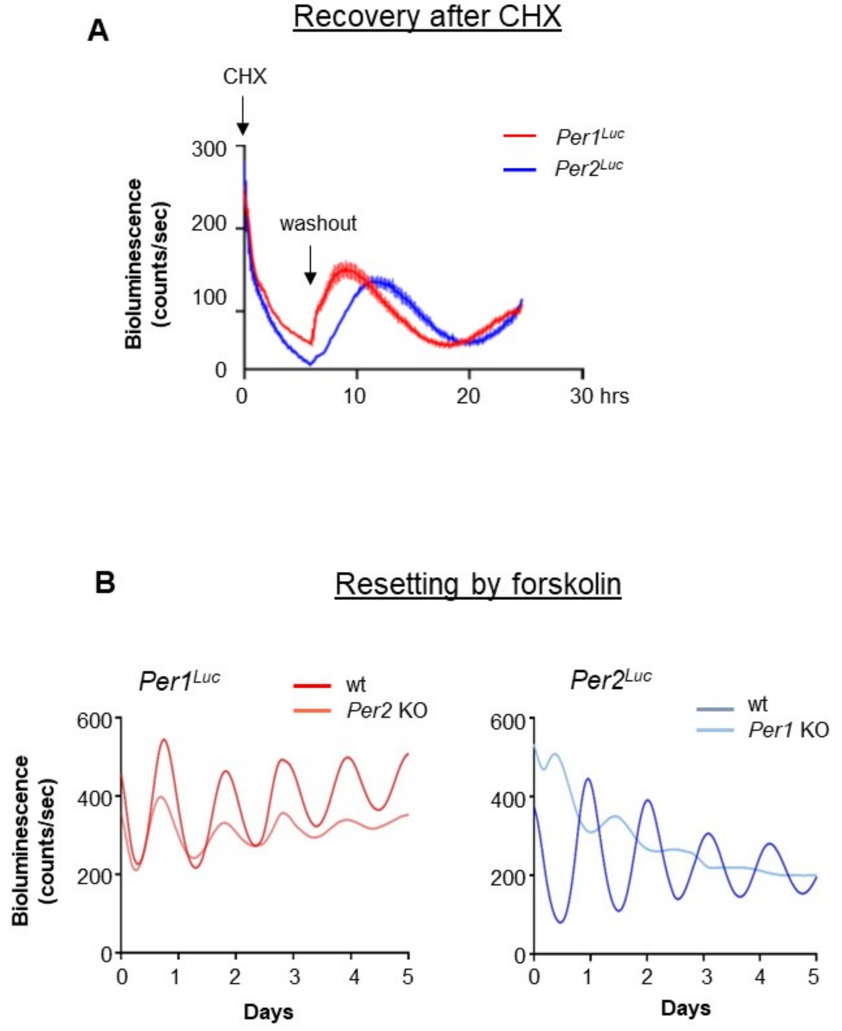


**Figure S5. PER1 and PER2 are different in key parameters.** (A) De novo PER1 accumulated faster than PER2. Existing PER was depleted by CHX treatment for 8 hrs. mean+/-SEM, n=3, $p<0.001$. (B) Antiphase oscillation of PER2-Luc in *Per1* KO cells was reproduced by forskolin treatment. Related to Figure 3 and 4.

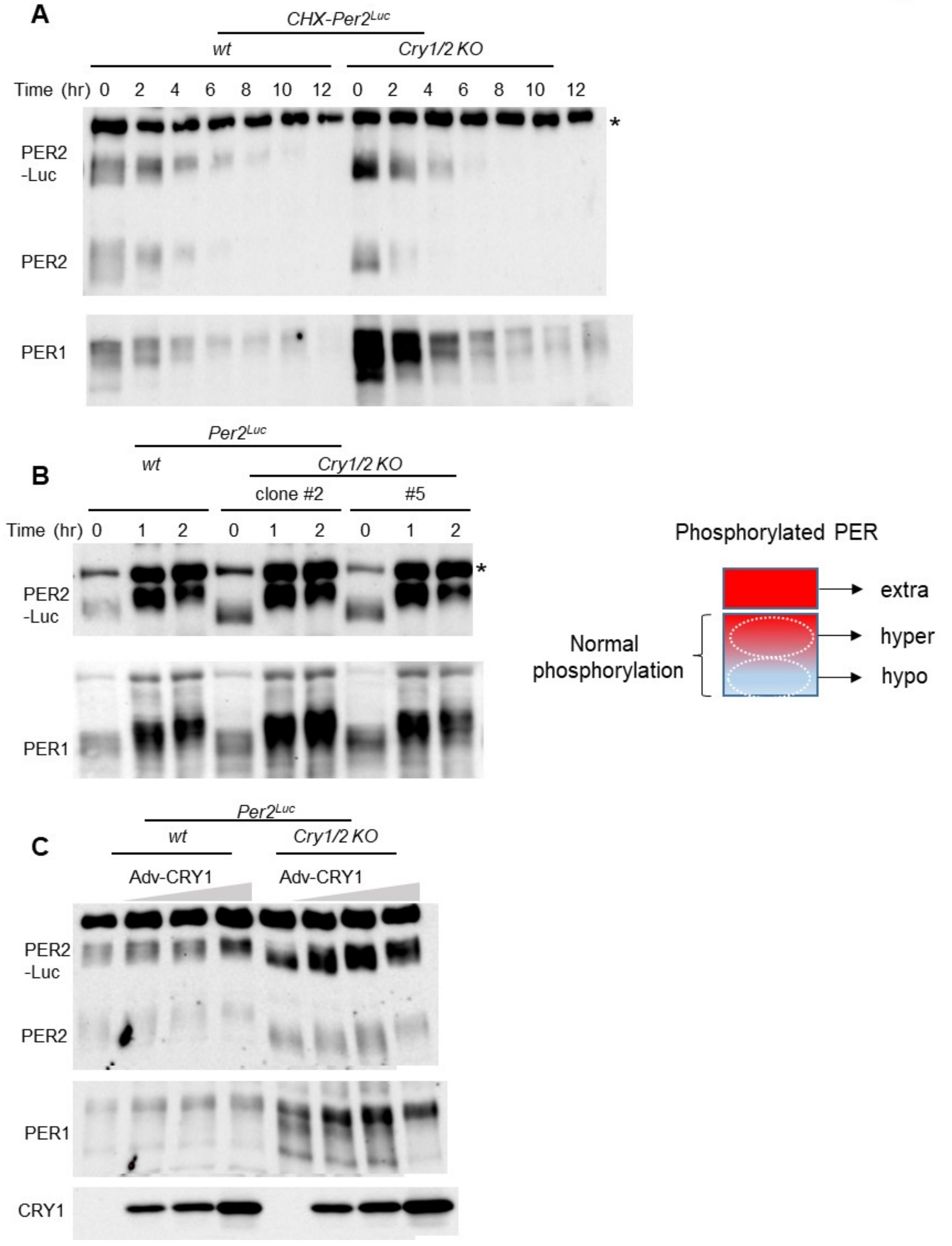


**Figure S6. CRY protects hyperphosphorylated PER from rapid degradation.** (A) PER proteins were degraded before reaching a hyperphosphorylated state in *Cry1/2* KO cells. (B) PER can be hyperphosphorylated in *Cry1/2* KO cells as much as PER in wt cells. This suggests that hyperphosphorylated PER species are unstable without CRY. Three stages of PER phosphorylation are illustrated in the left panel. Extra hyperphosphorylated species are not detected in a steady state because they are very unstable but can be detected in *β-Trcp1/2* KO cells (D'Alessandro et al., 2017). (C) Hyperphosphorylated PER species accumulated in *Cry1/2* KO cells when transgenic CRY1 was expressed. Related to Figure 5.

Fig S7

```
                                 E2
hCK1e          ---------MELRVGNKYRLGRKIGSGSFGDIYLGANIASGEEVAIKLECVKTKHPQLHIE
hCK1d          ---------MELRVGNRYRLGRKIGSGSFGDIYLGTDIAAGEEVAIKLECVKTKHPQLHIE
hCK1a          MASSSGSKAEFIVGGKYKLVRKIGSGSFGDIYLAINITNGEEVAVKLESQKARHPQLLYE
                         *: **.:*:* *************. :*: *****:***. *::****  *

hCK1e          SKFYKMMQGGVGIPSIKWCGAEGDYNVMVMELLGPSLEDLFNFCSRKFSLKTVLLLADQM
hCK1d          SKIYKMMQGGVGIPTIRWCGAEGDYNVMVMELLGPSLEDLFNFCSRKFSLKTVLLLADQM
hCK1a          SKLYKILQGGVGIPHIRWYGQEKDYNVLVMDLLGPSLEDLFNFCSRRFTMKTVLMLADQM
               **:**::******* *:* * * ****:**:***************:*::****:*****

hCK1e          ISRIEYIHSKNFIHRDVKPDNFLMGLGKKGNLVYIIDFGLAKKYRDARTHQHIPYRENKN
hCK1d          ISRIEYIHSKNFIHRDVKPDNFLMGLGKKGNLVYIIDFGLAKKYRDARTHQHIPYRENKN
hCK1a          ISRIEYVHTKNFIHRDIKPDNFLMGIGRHCNKLFLIDFGLAKKYRDNRTRQHIPYREDKN
               ******:*:*******:********:*:: * :::*********** **:*******:**
              E4
hCK1e          LTGTARYASINTHLGIEQSRRDDLESLGYVLMYFNLGSLPWQGLKAATKRQKYERISEKK
hCK1d          LTGTARYASINTHLGIEQSRRDDLESLGYVLMYFNLGSLPWQGLKAATKRQKYERISEKK
hCK1a          LTGTARYASINAHLGIEQSRRDDMESLGYVLMYFNRTSLPWQGLKAATKKQKYEKISEKK
               ***********:***********:***********  ************:****:*****

hCK1e          MSTPIEVLCKGYPSEFSTYLNFCRSLRFDDKPDYSYLRQLFRNLFHRQGFSYDYVFDWNM
hCK1d          MSTPIEVLCKGYPSEFATYLNFCRSLRFDDKPDYSYLRQLFRNLFHRQGFSYDYVFDWNM
hCK1a          MSTPVEVLCKGFPAEFAMYLNYCRGLRFEEAPDYMYLRQLFRILFRTLNHQYDYTFDWTM
               ****:******:*:**: ***:**.***:: *** ******* **:   ...***.***.*
                                        E8
hCK1e          LKFGAARNPEDVDRERREHEREERMGQLRGSATRALPPGPPTGATANRLRSAAEPVASTP
hCK1d          LKFGASRAADDAERERR--DREERLRHSRNPATRGLPSTASG-----RLRGTQEVAPPTP
hCK1a          LKQKAAQQAASSSGQGQ-------------------------------------------
               **  *:: . . . : :
                    Per1Luc CKKO-2: REE → - - P

hCK1e          ASRIQPAGNTSPRAISRVDRERKVSMRLHRGAPANVSSSDLTGRQEVSRIPASQTSVPFD
hCK1d          LTPTSHTANTSPRPVSGMERERKVSMRLHRGAPVNISSSDLTGRQDTSRMSTSQIPGRVA
hCK1a          -------------------------QAQTPTGKQTDKTKSNMKGF----------------
                                        . :   *  .: :.*::.*

hCK1e          HLGK------
hCK1d          SSGLQSVVHR
hCK1a          ----------
```

C-terminus alignment

```
hCK1e          FGAARNPEDVDRERREHEREERMGQLRGSATRALPPGPPTGATANRLRSAAEPVASTPAS
hCK1d          FGASRAADDAERERR--DREERLRHSRNPATRGLP-----STASGRLRGTQEVAPPTPLT
               ***:* .:*.:****  :****: : *..***.**     .:::.***.: * ...** :

hCK1e          RIQPAGNTSPRAISRVDRERKVSMRLHRGAPANVSSSDLTGRQEVSRIPASQTSVPFDHL
hCK1d          PTSHTANTSPRPVSGMERERKVSMRLHRGAPVNISSSDLTGRQDTSRMSTSQIPGRVASS
                 . :.*****.:* ::**************.*:*********:.**:.:** .  .

hCK1e          GK------
hCK1d          GLQSVVHR
               *
```

**Figure S7. Catalytic domains of CK1 isoforms are well conserved.** Three most conserved human CK1 isoforms are aligned. CRISPR target sites are represented by red arrows; E2 and E8 for CK1ε, E4 for CK1δ. C-termini are highlighted in yellow. The red boxed sequence was mutated from Arg Glu Glu to del del Pro in the *Per1*$^{Luc}$-CKKO-2 clone. Related to Figure 6.

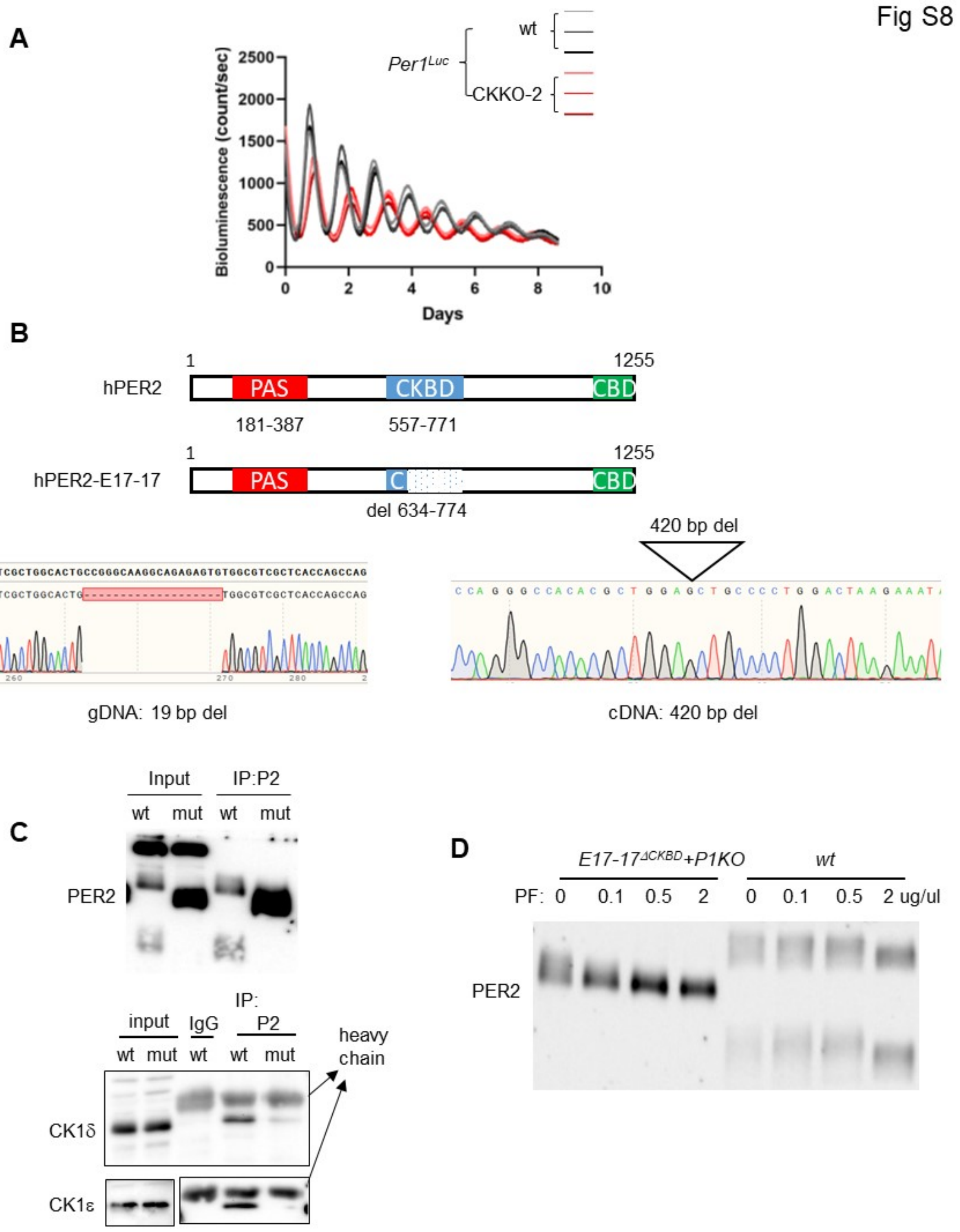


**Figure S8. Robust interaction between PER-CKBD and CK1δ/ε is critical for the phosphotimer.** (A) *Per1*$^{Luc}$-CKKO-2 clone showed a ~28.5 hr period which is comparable to other *CK1* double mutant clones and much longer than *CK1δ* or *ε* single mutant clones. Another bioluminescence rhythm plot is shown. Three replicates of wt and the mutant are shown. (B, C) A PER2$^{\Delta CKBD}$ mutant is defective in binding to PER. One allele had a frame-shifting mutation while the other allele had a large deletion. Sequencing of cDNA revealed that the deletion-mutant allele had an in-frame deletion missing 2/3 of CKBD. CoIP is representative of two experiments. (C) Hyperphosphorylation of the mutant PER2 was more sensitive to the CK1δ/ε inhibitor. Related to Figure 6 and 7.

Fig S9

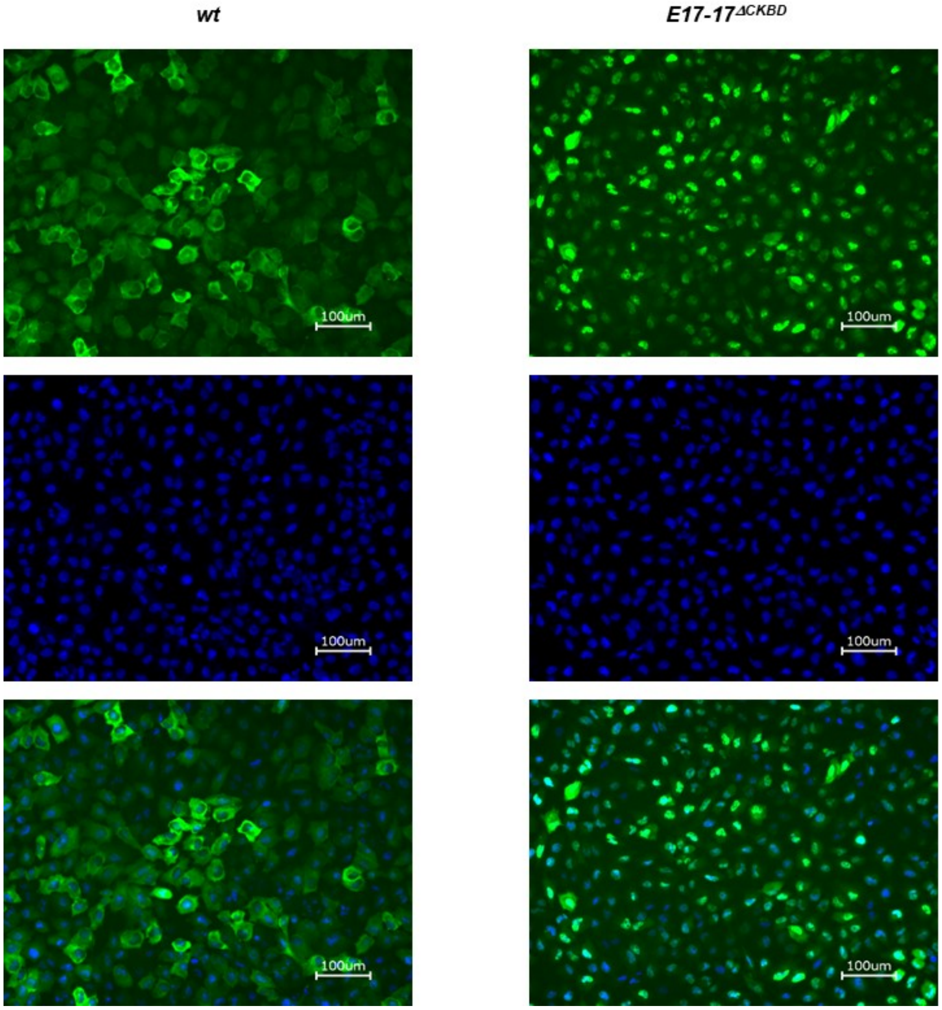


**Figure S9. PER2$^{\Delta CKBD}$ is predominantly nuclear.** PER2-Venus and PER2$^{\Delta CKBD}$-Venus fusion proteins were expressed using the adenoviral vector as we have done previously (Beesley et al., 2020). Venus fluorescence was visualized using confocal microscopy. Related to Figure 7.
